# CX3CR1 And MHCII Define Distinct Synovial Macrophage Populations With Conserved Inflammatory Response Across Mouse Models Of Rheumatoid Arthritis

**DOI:** 10.64898/2026.07.30.741914

**Authors:** Yidan Wang, Shang-Yang Chen, Philip Homan, Anna Montgomery, Jessica Maciuch, Sam D. Dowling, Salina Dominguez, Gaurav Gadhvi, Tyler A. Therron, Blair Eckman, Ana Teodósio, Dawn Howdle, Vanessa Manada De Lobos, Mohammad D. Khan, Kainat Mian, Meghan Lynn Mayer, Miranda G. Gurra, Hadijat-Kubura Moradeke Makinde, Matt Dapas, Andrew Filer, Carla Marie Cuda, Harris Perlman, Deborah R. Winter

## Abstract

Macrophages in the synovial lining are critical for the maintenance of healthy tissue while also contributing to the pathogenesis of rheumatoid arthritis (RA). However, the field currently lacks a unifying characterization of synovial macrophage heterogeneity across steady-state and inflammation. Here, we defined 4 transcriptionally distinct populations of synovial macrophages CX3CR1^+^MHCII^-^ (lining); CX3CR1^-^MHCII^-^ (interstitial/sublining); CX3CR1^-^MHCII^+^ (monocyte-derived); and CX3CR1^+^MHCII^+^ (infiltrating). The MHCII^-^populations are long-lived and derived from embryonic precursors regardless of localization while the MHCII^+^ populations differentiate from bone marrow (BM) progenitors dependent on CCR2. We identified conserved activation pathways between acute and chronic mouse models of inflammatory arthritis as well as novel arthritis-associated subpopulations. During peak inflammation, the influx of BM-derived cells was associated with upregulation of monocyte-related genes with concurrent down-regulation of tissue-resident genes. Our results provide a unifying schema for describing synovial macrophages across conditions and pave the way for future studies in modulating transcriptional activity in rheumatoid arthritis.

## Introduction

Rheumatoid arthritis (RA) is a chronic inflammatory and destructive arthropathy of unknown etiology. Previous studies have shown that the number of sublining macrophages relates to disease activity, progression of radiographic joint destruction and is an independent risk factor for severe progression of bone erosion [1–3]. Further changes in the numbers of sublining macrophages are currently the only known biomarker that is associated with response to therapy [4, 5]. We are also the first to show that a subpopulation of synovial macrophages (CD206^+^) in RA patients variably express 6 different transcriptional modules that are associated with medication status and joint disease severity [6].

The accelerating medicine partnership (AMP) was the first to utilize single cell RNA-seq (scRNA seq) on isolated CD14^+^ cells from the synovium of RA and osteoarthritis (OA) patients. AMP described 4 different populations of synovial macrophages which, based on abundance in different patients and enriched GO processes, were thought to differentially contribute to RA or OA [7]. Huang et al [8] and Ng et al [9] also identified 4 population of macrophages from OA patients undergoing total knee arthroscopy and from frozen shoulder. A more extensive manuscript with a larger dataset was published by AMP detailing 15 subpopulations of myeloid cells including 9 macrophage populations [10]. The authors identified cell-type abundance phenotypes (CTAPs) that are enriched in specific cell states, although CTAPs did not correlate with clinical outcomes [10]. Additional studies were able to separate populations based the status of anticitrullinated protein antibody. Wu et al demonstrated that the proportion of HLA-DRB5^hi^CD36^+^ were lower, while HLA-DRB5^-^CCL^+^ and IL18^+^CCL^+^ synovial macrophages were increased in APCA^-^ vs ACPA^+^ RA patients [11]. Jiang et al showed that C1Q^hi^ and MARCO^hi^ were higher in ACPA^-^ and APOE^hi^ and FN1^hi^ synovial macrophages were enriched in ACPA^+^ RA patients [12]. Alivernini et al identified 9 different subpopulations of myeloid cells in the synovium of healthy controls and RA patients [13] using scRNA seq. These were aligned with the four major subtypes of macrophages distinguishable by flow cytometry. Based on their data, the authors compared the proportions of subpopulations across different classes of RA (naïve, resistant, remission) and healthy controls. Hanlon et al [14] reported similar results to Alivernini et al including 9 subpopulations of macrophages based on flow cytometry and scRNA seq. More recently, Mantel et al identified one of the macrophage sublining populations that express SPP1 colocalized with fibrin [15].

While early research in the 1990s postulated that the synovial lining may be comprised of a heterogenous population of fibroblasts and macrophages, actual markers that distinguish their phenotypes or functions were not yet realized. We are one of the first to demonstrate heterogeneity in the mouse synovial macrophage compartment [16]. We identified at least two populations of synovial macrophages based on the expression of MHCII that differ in their origin, radio-sensitivity, requirement of MCSF for survival, expression of genes known to be involved in inflammation, repair and macrophage development and function in the synovium during steady state and inflammatory arthritis [16]. These macrophages are similar to those observed in the heart [17], kidney [18], and liver [18] and represent tissue-resident macrophages and bone marrow-derived macrophages, respectively. Culemann et al is the first to reveal the topology of the individual subpopulations of synovial macrophages using confocal immunofluorescent microscopy and 3D-light-sheet fluorescence microscopy of optically cleared mouse knee joints [19]. The authors show that CX3CR1^+^ synovial lining cells develop embryonically, have a half-life of 5 weeks and do not require recruited monocytes to maintain homeostasis, but may be replenished by MHCII^+^CSF1R^+^CX3CR1^-^interstitial synovial macrophages, similar to our previous studies [16]. Culemann et al also identified 6 subpopulations of murine macrophages including CX3CR1^+^ lining cells, AQP1^+^, MHCII^+^, and RELM-a^+^ interstitial macrophages, ACP5^+^ osteoclast precursors, and ^i^proliferating cells on FACsorted CD45^+^CD11b^+^Ly6G^-^ synovial cells at steady-state and in early stages of serum transfer induced arthritis (STIA). While functional studies have since been performed that assign functions to different macrophage populations based on specific markers, it is not entirely clear how these align to previously defined subsets [20, 21]. Moreover, other groups have categorized the macrophage subpopulations in the mouse synovium differently than Culemann et al during homeostasis as well as during anti-GPI induced inflammatory arthritis [22] or post-traumatic osteoarthritis [23]. In our published study in the aging joint, we were also unable to fully recapitulate the populations described [24]. These data demonstrate that there is an unmet need to classify the heterogenous macrophage population in multiple models of inflammatory arthritis.

In this study, we employed an integrative approach of immunological and genomic technologies to define four distinct major populations of synovial macrophages. We assessed the ontogeny and turnover of these populations at steady-state using parabiosis, bone marrow chimeras, fate-mapping, and genetic knockouts. We then characterize these populations across three models of inflammatory arthritis: STIA, collagen-induced arthritis (CIA), and KRNAg7 spontaneous arthritis (KRN). Through FACS followed by population level RNA-seq, we investigate up and down-regulated pathways in acute (STIA) and chronic (CIA) inflammation. Using single-cell transcriptional profiling, we assess changes in synovial macrophage heterogeneity and origin in the inflamed joint. Our results demonstrate the consistency of the major synovial macrophage populations across models and defined conserved activation pathways in response to joint inflammation.

## Materials and Methods

### Mice

All mice used in the study were females (with the exception of the male DBA/1 mice in the CIA model) and aged between 8-12 weeks or 8-20 weeks for post-bone marrow restoration of bone marrow chimeras (BMC) mice. All strains of mice were on C56BL/6 (B6) background. Wild-type (B6.CD45.2, JAX: 000664), DBA/1, B6.CD45.1 (JAX: 033076), KRN, Ag7, KRNAg7, NOD, CCR2^-/-^(JAX: 004999), CX3CR1^ERCre/+^, and zsGFP mice were bred and housed in the specific pathogen-free, barrier animal facility of Northwestern University. All experimental procedures on mice reported in the study were approved by Institutional Animal Care and Use Committee (IACUC) office.

### K/BxN serum transfer induced arthritis (STIA)

The generation of STIA mice was as previously described by this group [16, 25, 26]. Serum from K/BxN transgenic mice was collected and administered retro-orbitally to mice by assessing ankle swelling using clinical scores every 2–4 days using the following scoring system: 0 = no inflammation; 1 = swollen toes with the ankle maintaining a funnel shape; 2 = swollen ankle with the funnel shape disappearing; 3 = swollen ankle with an inverted funnel shape. Independent titrated K/BxN serum was used to induce STIA for each experiment.

### Collagen-Induced Arthritis (CIA)

Five milligrams of type II bovine collagen (lyophilized) were dissolved in 2.5mL of 0.01M (10mM) Acetic acid and left constantly rolling at 4°C overnight. Complete Freud’s Adjuvant (CFA) was vortexed well before adding to collagen/acetic acid solution. Prepared collagen was combined in a 1:1 ratio with CFA (2.5mL CFA to 2.5mL of collagen in acetic acid) drop-wise as the emulsion mixed using a magnetic stirring bar until a viscous emulation was produced. Mice were anesthetized and a single intradermal injection of 100µL of the emulsion was administered approximately 0.25 cm above the base of the tail. Booster immunizations (day 21) are administered using same method as day 0 (Emulsion preparation and administration) with incomplete Freud’s Adjuvant (IFA) instead of Complete Freud’s Adjuvant (CFA). Mice were scored every 4-6 days until day 62 after immunization using the extended scoring system (0-60 per mouse, 0-15 per paw).

### KRNAg7 Spontaneous Arthritis

The generation of KRNAg7 was generated as previously described [27]. Male KRN mice on C57BL/6 background were crossed with female Ag7 congenic mice on C57BL/6 background. Spontaneous arthritis occurs in KRNAg7 mice starting at 4-week-old and persists throughout the life. Ankles were collected for single-cell RNA-seq (scRNA-seq) from 8-12 week-old KRNAg7 mice.

### Fate-Mapping Studies

Studies using CX3CR1^ERCre/+^zsGFP mice were carried out as previously described (Yona et al, 2013). To establish embryonic origins, 50mg/kg tamoxifen (Sigma) and 10mg/mL progesterone (Sigma) was dissolved in corn oil and administered by oral gavage at either E8 or E15 of gestation to pregnant CX3CR1^CreER^zsGFP mice. The pups were then sacrificed at adulthood (approximately 8 weeks) to determine the percent of macrophages that were GFP positive. As a comparison, 50mg/kg tamoxifen in corn oil was administered intraperitoneally (i.p.) on consecutive days to adult mice, which were sacrificed 1 day later (Tam positive) . Non-tamoxifen controls were also included to reflect baseline GFP expression (Naïve). To assess turnover at steady-state and in inflammation, tamoxifen was administered on two consecutive days to adult mice who were then either subjected to STIA or untreated as controls. Mice were euthanized at various time points following the second dose of tamoxifen. In addition, separate cohorts of mice were treated with tamoxifen on Day 3 following STIA induction and euthanized on following time points.

### Bone marrow chimeras

Bone marrow chimera (BMC) mice were generated as previously described [25, 26]. Antibiotic-treated water containing 1mL of Sulfamethoxazole and Trimethoprim (Novitium Pharma, 70954-258-10) per 250 mL drinking water was provided to recipient mice, including B6.CD45.1, B6.CD45.2, and Ccr2^-/-^ mice, for a total of 6 weeks: 1 week before and 5 weeks after BMC generation. Prior to irradiation the anesthesia treatment of 100 microliters (µL) containing 100 milligram (mg)/1 kilogram (kg) mouse weight Ketamine (Dechra, ANADA 200-073) and 20mg/kg xylazine (AnaSed Injection, NDC 59399-110-20) diluted in sterile water were administered intraperitoneally to the recipient mice. All ankles joints were shielded with lead during irradiation to preserve the synovial macrophage compartment [25, 26]. Irradiation was performed at 100 centigray (cGy) γ irradiation on each host mouse. To completely ablate remaining bone marrow stem cells, 40mg/kg busulfan (Caymen Chemical, 55-98-1) was administered intraperitoneally (IP) to the recipient mice 6 hours post the irradiation procedure. One day post the irradiation procedure, a minimum of 10 million cells from the bone marrow of donor mice with different CD45 allotype, was administered retro-orbitally to the recipient mice. Peripheral blood (PB) from the submandibular vein was used to determine the success of chimerism 6-8 weeks post bone marrow cell transfer.

### Parabiosis

The generation of parabiotic mice was performed at Microsurgery Core, Northwestern University as described in Kamaran P, *et al.* [28] . Antibiotic-treated water containing 1 mL of Sulfamethoxazole and Trimethoprim (Novitium Pharma, 70954-258-10) per 250 mL drinking water was provided to mice for a total of 6 weeks, including 1 week before the surgery and 5 weeks after the surgeries. To keep parabiotic mice hydrated, daily subcutaneous sterile saline (Teknova, S5820) was administrated to mice subcutaneously for a total of 2 weeks following surgery.

### Clodronate laden liposome (Clo-lip) administration

A total volume of 200 µL of Clo-lip (LIPOSOMA, 5mL) was administered retro-orbitally to BMC mice. The numbers of circulating monocytes in peripheral blood and synovial macrophages in joint tissue were assessed by flow cytometry on days 0, 1, 3, 7 post Clo-lip administration. A separate cohort of BMC mice treated with Clo-lip was collected for CITE seq at day7 post administration.

### Preparation of Synovial Cells

Isolation of synovial cells was performed as previously described [25, 26]. Joints were removed from hindpaw in pairs following euthanasia and perfusion. Skin and toes were removed from each paw and bone marrow was flushed from exposed tibia with sterile 1 x HBSS. Synovial tissue was then infused with 1.5mL/joint of ankle digestion buffer (2.4mg/mL dispase II, 2mg/mL collagenase D, 0.2mg/mL DNAse I in HBSS pH 7.2-7.6) before incubation at 37C for 1h with shaking. Cells were then agitated through a 40 μm mesh filter. Erythrocytes lysis was performed using 200 μL 1x PharmLyse for 1 minutes at room temperature (BD Biosciences, 555899).

### Flow cytometry analysis

Cardiac puncture or submandibular vein methods were used for PB collection in EDTA K3E 1.3 mL microsample tube (Sarstedt Inc, 41.1395.105) as previously described [16, 25]. PB single cell suspension of 90μL was incubated in 0.5mg CD16/32 Fc Block (BD Biosciences, 553142) for 20 minutes at 4ß to prevent non-specific binding to Fc receptors on immune cells. Following Fc block incubation, the PB single cell suspension was stained with an antibody cocktail mix for 30 minutes at 4ß. The antibody mix includes R718 CD45.1 (BD Biosciences, 567297), BV421 CD45.2 (BioLegend, 109831), BV711 Ly6G (BD Biosciences, 563979), PE-CF594 SiglecF (BD Biosciences, 562757), BB700 CD11b (BD Biosciences, 566416), APC CD4 (BD Biosciences, 553051), APC CD8a (BD Biosciences, 553035), APC CD19 (BD Biosciences, 550992), APC NK1.1 (BD Biosciences, 550627), PE CD115 (Invitrogen, 12-1152-82), BV650 CX3CR1 (Biolegend, 149033), PE Cy7 CD62L (BioLegend, 104418), APC Cy7 (BD Biosciences, 560596). After staining, 1mL FACS Lyse (BD Biosciences, 349202) was used for red blood cell lysis and fixation at room temperature for 10 minutes. 123count eBeads (Thermo Fisher, 01-1234-42) were added for absolute cell count calculation, after which the samples were ready for data acquisition. Dead cells were stained with eFluor 506 viability dye (eBioscience) (1:1000 dilution). Cells were incubated with FcBlock (BD Bioscience) and stained with antibodies (see table) for flow cytometry analysis and fluorescence-activated cell sorting (FACS). 123count eBeads (Thermo Fisher, 01-1234-42) were utilized to calculate cell counts. A 2% of paraformaldehyde solution (Electron Microscopy Sciences, 15713-S) diluted in 1 x PBS was used for fixation at room temperature in dark for 15 minutes. All flow cytometry analytical data were obtained on LSR II, Symphony A5 Analyzer, and Symphony A5.2 Spectral Analyzer (BD Biosciences) at Robert H. Lurie Comprehensive Cancer Center (RHLCCC) Flow Cytometry Core Facility,

Northwestern University. FACSAria^TM^ III Cell Sorters (BD Biosciences) configured with 4 or 5 lasers were used for cell sorting. When required, gating boundaries were determined by fluorescence minus one (FMO). FlowJo (v10.7.1) was used for compensation calculation, spectral unmixing and downstream analysis.

### Processing of Single-Cell RNA Sequencing Libraries

For all single-cell studies, live CD45^+^CD11b^+^Ly6G^-^SiglecF^-^CD64^+^ synovial macrophages from ankle joints were sorted from a single-cell suspension. Generation of GEMs, barcoding, cDNA amplification, 3′ gene libraries, and cell surface protein libraries was conducted using the Chromium Next GEM Single Cell 3′ (v3.1, dual index) protocol at the Metabolomics Core, Northwestern University. For datasets stained with Antibody-Derived Tags (ADT) – Steady State (SS), BMC Clo-Lip, BMC STIA, a Feature Barcode library was generated following v3.1 step 4 cell surface protein library construction in parallel with gene expression (GEX) libraries). Sequencing was performed on a NextSeq 2000 (Illumina) at the same facility.

### Quality Control of Single-Cell Datasets

The mouse reference genome mm10 (mm10-2020-A) was used for read alignment and quantification through the count function of the Cell Ranger pipeline (v6.1.2). Seurat (v5.0.3) package was utilized for quality control, preprocessing, clustering, identification of *de novo* markers, label transfer, and visualization. Quality control (QC) for each dataset was assessed based on UMI counts, mitochondrial gene percentages, and ADT counts thresholds were set as defined below. Doublets were identified and removed using scDblFinder (v1.12.0) with an expected doublet formation rate of 5% [29]. Pre-processing and analysis were performed using Seurat V5 in R 4.3.0 for all single-cell data sets [30, 31]. Data sets with multiple samples were merged, but all pre-processing steps until dimensional reduction were performed separately by sample. Data sets were filtered to remove low-quality cells according to the parameters in **Table 1**. Thresholds were determined using a post-hoc filtering method in which cells were clustered prior to QC filtering and thresholds were set to exclude clusters with abnormal QC and a lack of distinct marker genes. Cells with ADT UMI count above the threshold were excluded from ADT-specific analyses and visualizations but not filtered from the dataset.

**Table 1.** Single Cell Data Set Pre-processing Parameters.

| **Dataset** | **# cells**<br>**pre-QC** | **#doublets**<br>**removed** | **RNA UMI**<br>**Count** | **% Mito-**<br>**chondrial** | **ADT UMI**<br>**Count** | **# cells**<br>**post QC** |
| --- | --- | --- | --- | --- | --- | --- |
| **SS**<br>**CITE-seq** | 9914 | 1030 | > 6000 | < 7.5 | <15000 | 5376 |
| **BMC**<br>**Clo-lip**<br>**CITE-seq** | Ctrl: 6296<br>D7: 5622 | Ctrl: 585<br>D7: 509 | > 3000 | < 10 | < 7500 | Ctrl: 3034<br>D7: 3128 |
| **BMC**<br>**STIA**<br>**CITE-seq** | D0: 9545<br>D1: 6511<br>D7: 7053<br>D14: 10561<br>D56: 11142 | D0: 976<br>D1: 634<br>D7: 716<br>D14: 1044<br>D56: 1033 | >3000<br>< 60000 | < 10 | < 7500 | D0: 5838<br>D1: 4122<br>D7: 4141<br>D14: 4633<br>D56: 7076 |
| **KRN** | 3765 | 296 | >1500<br>< 20000 | <10 | NA | 2794* |

\*For KRNag7 an additional filter of % ribosomal RNA <40 was applied

### Clustering of Single-Cell Datasets

RNA counts were separately normalized using Seurat’s SCTransform function for use in dimension reduction and clustering. Module scores were calculated for S and G2M phase cycling gene lists provided by Seurat (translated into mouse orthologs using biomaRt [32, 33] and regressed out during SCTransform normalization. ADT assays were normalized using the centered log ratio method within each cell (margin = 2). Variable genes (n=750) were used for PCA dimensional reduction and clustering was performed using Seurat’s Leiden algorithm implementation on a shared nearest neighbor graph with the parameters specified in **Table 2**. Clustering parameters were chosen by minimizing the number of PCs and clustering resolution that allowed for separate clustering of known populations of interest. De-novo cluster markers were identified using the FindAllMarkers function in Seurat on the log-normalized RNA assay (assay = “RNA”), restricting top markers to those expressed in at least 50% of cells within each cluster (min.pct = 0.5).

**Table 2.** Single Cell Data Set Clustering Parameters.

| **Dataset** | **# PCs** | **Clustering Resolution** | **# de novo clusters** |
| --- | --- | --- | --- |
| SS CITE-seq | 16 | 0.6 | 6 |
| BMC Clo-lip CITE-seq | 11 | N/A | N/A |
| BMC STIA CITE-seq | 10 | 0.2 | 8 |
| KRN | 13 | 0.18 | 8 |

### Label Transfer of Population Annotations

For each dataset after Figure 1, *de novo* clusters annotated as macrophages were subset out.

**Figure 1.**
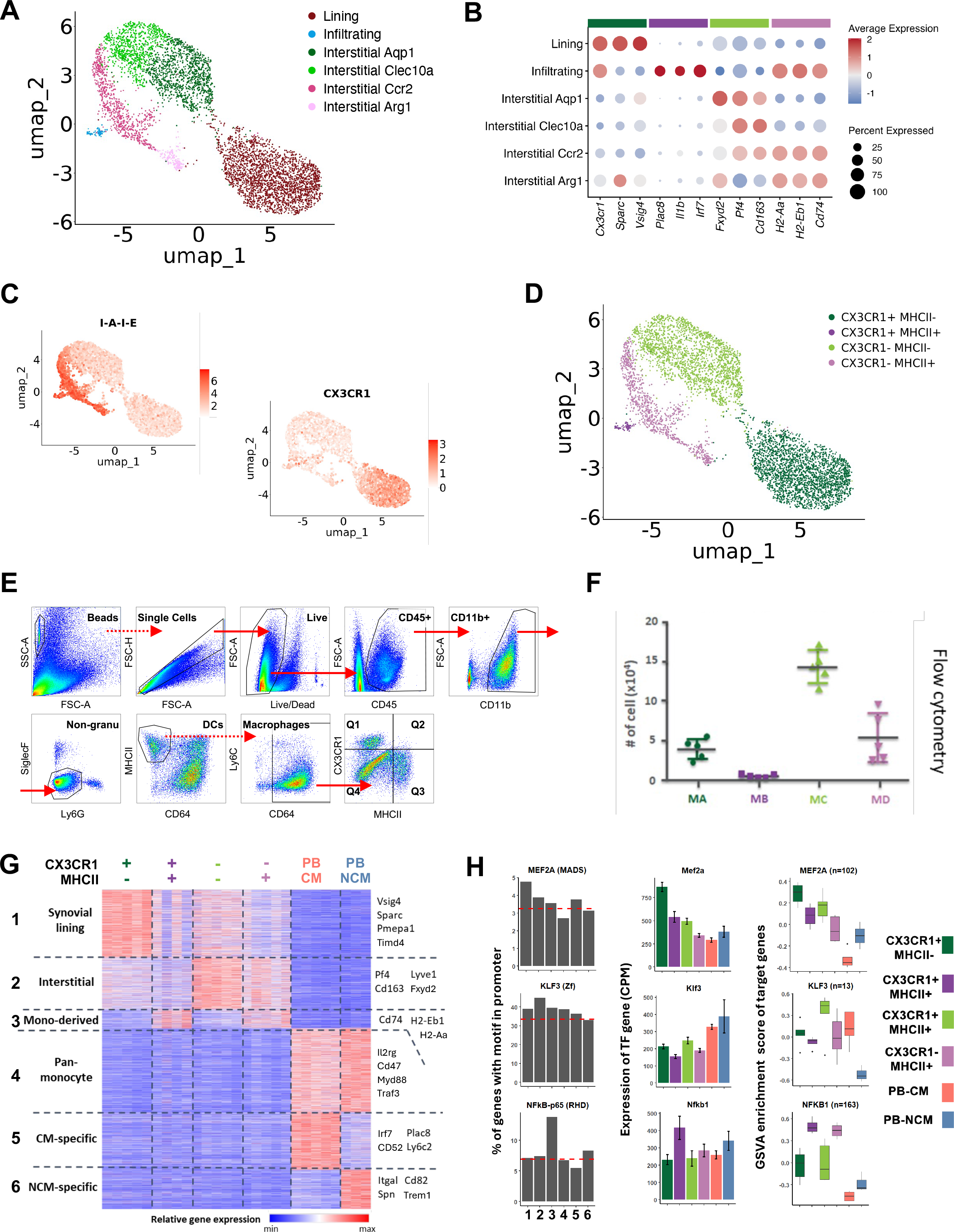
CITE-seq identified 4 synovial macrophages align well with bulk RNA-seq defined 4 synovial macrophages. **(A)** Uniform Manifold Approximation and Projection (UMAP) depicting 6 functionally distinct synovial macrophage subpopulations identified by Cellular Indexing of Transcriptomes and Epitopes by Sequencing (CITE-seq) of liveCD45^+^CD11b^+^CD4^-^CD8^-^CD19^-^Ly6G^-^SiglecF^-^CD64^+^ synovial cells from C57BL/6 (B6) mice. **(B)** Bubble plot showing selected key genes of 6 synovial macrophage subpopulations. **(C)** Feature plots of antibody derived tag (ADT) CX3CR1 and I-A-I-E (MHCII) Expression. **(D)** UMAP depicting 4 distinct synovial macrophage subpopulations based on the ADT expression of CX3CR1 and MHCII. **(E)** Flow cytometry gating strategy of 4-macrophage subpopulation distinguished by CX3CR1 and MHCII surface antibodies. **(F)** Absolute numbers of 4 macrophage subpopulations identified by flow cytometry. **(G)** Gene patterns identified by k-means clustering of the bulk RNA-seq results of 4 macrophage subpopulations and peripheral blood (PB) classical monocytes (CM) and non-classical monocytes (NCM).

These cells were further annotated by label transfer using the SS CITE-seq dataset as reference. FindTransferAnchors and TransferData functions from Seurat were used. Transferred labels were merged with de novo annotation of non-macrophages to generate the entire dataset annotation.

### Additional Analyses of Single-Cell Data

Similarity between *de novo* clusters or populations across datasets was calculated by Pearson correlation between the log-normalized average cell gene expression of the union of variable genes in datasets. SingleR was used to calculate similarity of individual cells to a custom bulk RNA-seq reference created by averaging CPM-normalized values across replicates for each steady-state population [34]. For the BMC analyses, cells in the control sample were assigned a CD45 label based on their expression of CLR-normalized ADT values for CD45.1 and CD45.2 relative to a dataset-specific threshold based on non-immune CD45- cells. Gene modules for the BMC STIA dataset were generated using singleCellHaystack [35, 36] on the top 750 variable genes across all timepoints ranked by Kullback-Leibler Divergence. Genes were hierarchically clustered and the dendrogram was cut to yield 8 clusters. Module scores were then calculated for each cell using Seurat’s AddModuleScore function. Pathway enrichment for modules was performed using the enrichGO function from clusterProfiler [37], setting the background universe as all genes detected in at least 1% of cells in any sample.

Only pathways that passed a q-value threshold of 0.05 were retained. All correlation and pseudobulk heatmaps were visualized using ComplexHeatmap [38]. Bubble plots were generated by Seurat’s DotPlot with default settings. The CLR-normalized ADT values were averaged for and displayed as a heatmap.

### Bulk RNA-seq library preparation and analysis

Ten thousand to one hundred thousand cells were isolated by FACS for each of the four macrophage populations. RNA was extracted using PicoPure RNA Isolation kit (ThermoFisher) as per manufacturer’s instructions. Library prep was performed using QuantSeq 3’ mRNA sequencing kit (Lexogen) and sequenced on Illumina NextSeq. The resulting BCL sequencing files were demultiplexed using bcl2fastq into FASTQ format. The reads were trimmed using BBDuk v37.22 ( http://jgi.doe.gov/data-and-tools/bb-tools/), aligned to mm10 genome with STAR [39], and mapped to reference gene exons using HTseq [40] to generate a matrix of gene expression counts. Raw counts are normalized to counts per million (CPM) to account for differing read depth across samples. Expressed genes were defined as those with expression greater than 16 CPM in at least 4 samples. This results in 7668 expressed genes for SS, 8513 genes for STIA, and 9388 genes for CIA datasets. Principal component analysis (PCA) was performed using the prcomp function with data scaling and centering. For calculating the expression fold-change relative to Day 0 of STIA and CIA timecourse experiments, CPM values lower than 16 were set to 16 to minimize signal from lowly expressed genes. Temporal differentially expressed genes (DEGs) across the STIA and CIA time courses were defined as those with at least 2-fold change in expression between day 0 and any subsequent time points in at least one macrophage population. DEGs were designated as acute, persistent, or resolution in STIA based on whether they were unique to Day 7, shared between Day 7 and Day 21, or unique to Day 21, respectively. Similarly, DEGs were designated as peak, chronic, or late in CA based on whether they were unique to Day 41, shared between Day 41 and Day 62, or unique to Day 62, respectively. K-means clustering and heatmap visualization on CPM values (SS) or log expression fold-changes relative to day 0 (STIA) of DEGs was performed using Morpheus web app (https://software.broadinstitute.org/morpheus). MAGNET webapp (https://magnet-winterlab.herokuapp.com/magnet[41]) was utilized to assess overlap between SS clusters against published gene sets from Lavin et al. [42] and Culemann et al [19]; between SS clusters and STIA clusters; and between CIA DEGs and STIA clusters. Gene ontology (GO) analyses of biological processes were performed using GOrilla webapp [43] with expressed genes as background. Transcription factor binding motif enrichment analysis was carried out with HOMER findMotifs.pl [44] using each k-means cluster as input and expressed genes as background. GSVA R package [45] was utilized to compute the combined relative expression scores for putative downstream target genes of select TFs. The downstream target genes for the select TFs were obtained from Dorothea database [46]by filtering for positive regulatory direction (mor=1) and using all levels of confidence (A-E). Unless otherwise stated, all computational analyses were performed using R v3.6.3, with figures generated via ggplot2 package.

### Sample preparation and staining for multiplex immunofluorescence

Sections were cut at 4 microns thickness, mounted on Superfrost Plus slides (VWR, 631-0448), dried at room temperature (RT) and baked overnight at 60°C prior to the staining. Slides were deparaffinized in two baths of xylene for 10 minutes each. Hydration was performed in a graded ethanol series (100%, 90%, 80%, 70%, 50%) and MilliQ water for 2 minutes each. Antigen retrieval was carried out on the Bond RX for 60 minutes using BOND Epitope Retrieval Solution 2. After retrieval, slides were kept at room temperature in Multistaining Buffer (MSB – Lunaphore, BU06) until loaded into the stainer. Staining was performed on the Lunaphore COMET platform (version 1.2.0.0). This system employs automated sequential immunofluorescence (seqIF™) staining, consisting of repeated cycles of primary antibody incubation, fluorescently labelled secondary antibody detection, imaging, and elution. All antibodies were tested and optimized for signal intensity and specificity, background noise, elution efficacy and epitope stability, using the software’s protocol templates (Screening, Optimization and Positioning). The full staining protocol was built based on the results of this workflow. Multiple staining cycles was performed per slide, one marker per cycle. Primary antibodies were incubated for 8 minutes at the dilutions specified and secondary antibodies were incubated for 4 minutes at a 1:400 dilution. Between staining cycles, elution was performed for 2 minutes at 37°C using Elution Buffer (Lunaphore, BU07-L). DAPI solution (Thermo Scientific, 62248) diluted at 1:1000 was used for nuclear counterstain.

### Statistical analysis

Statistical analysis of flow cytometry data was carried out in GraphPad Prism. P values were calculated using unpaired t test and were considered statistically significant if p<0.05.

## Results

### CX3CR1 and MHCII surface expression distinguish 4 distinct populations of synovial macrophages in mice

To investigate the heterogeneity of synovial macrophages in the murine synovium, we performed Cellular Indexing of Transcriptomes and Epitopes by sequencing (CITE-seq) on sorted CD45^+^CD11b^+^CD4^-^CD8^-^CD19^-^NK1.1^-^Ly6G^-^SiglecF^-^CD64^+^ cells from ankle joints of healthy mice. We then clustered the RNA-seq data to define 6 subpopulations which we annotated based on localization and key genes in previously published studies [19–21, 23, 24, 47] as lining, infiltrating, and interstitial (Aqp1, Clec10a, Ccr2, and Arg1-expressing) macrophages. (**Figure 1A, Supplemental Figure 1A-B**). However, these clusters are not robust and share common markers. Therefore, we used ADT levels to simplify these clusters into 4 major populations based on the expression of CX3CR1 and MHCII (**Figure 1B-D, Supplemental Figure 1C**): CX3CR1^+^MHCII^-^ synovial lining macrophages, CX3CR1^+^MHCII^+^ infiltrating macrophages, CX3CR1^-^MHCII^-^ interstitial macrophages, and CX3CR1^-^MHCII^+^ monocyte-derived macrophages. These annotations provide a more robust and consistent framework that can be used across flow cytometry and genomic studies.

To confirm our annotations, we sorted the four macrophage populations from ankle joints using FACS (**Figure 1E**). We found that the relative expression levels of key genes matched between single-cell and sorting-defined populations and that cells were generally assigned to the correct populations when the bulk RNA-seq data was used as reference (**Supplemental Figure 1D-E**). The number of cells in each population was highly consistent across mice and the transcriptional profiles were distinct from each other and robust across replicates (**Figure 1F, Supplemental Figure 1F**). We then utilized unsupervised k-means clustering to identify major gene expression patterns across the four macrophage populations, along with classical (CM) and non-classical (NCM) monocytes sorted from peripheral blood (PB) (**Figure 1G, Supplemental Figure 1G**). Monocyte samples were included in the clustering to better highlight gene signatures that are specific to synovial macrophages rather than constitutive in myeloid cells. We defined six expression patterns of which the first 3 were unique to the synovial macrophage populations. These macrophage-specific clusters all included genes associated with multicellular organismal process, biological quality, and cell differentiation as might be expected for any tissue macrophages. The first macrophage cluster, which was driven by CX3CR1^+^MHCII^-^ macrophages and to a lesser degree in CX3CR1^-^MHCII^-^, comprised genes that have been associated with synovial lining and tissue residency, such as Vsig4, Sparc, Pmepa1, and Timd4, as well as those involved in cellular homeostasis, cell adhesion, and ossification. This cluster overlaps the transcriptional signature seen in other long-lived, embryonic-derived tissue macrophage populations, such as microglia, Kupffer cells, and red pulp macrophages (**Supp Figure 1H**) [48]. The second macrophage cluster was shared between the CX3CR1^-^ populations, suggesting it reflects interstitial localization, and included previously reported markers of macrophage homeostasis, including Pf4, Cd163, Lyve1, and Fyxd2 [49–51]. The third macrophage cluster was highest in the MHCII^+^ populations and consists of genes associated with monocyte-derived origin such as those involved in cell chemotaxis, inflammatory response, and exogenous antigen presentation (e.g., Cd74, H2-Aa, H2-Eb1). This cluster is most enriched for genes shared between ileal and colonic macrophage populations which are known to have high replacement from monocytes (**Supp Figure 1H**) [52, 53]. Overall, these gene expression patterns were highly consistent with those observed in synovial lining and interstitial populations defined by Culemann et al (**Supp Figure 1H**) [19].

Finally, to determine the transcription factors (TFs) that might be driving macrophage expression patterns, we performed motif analysis to identify known TF motifs enriched in promoters of genes within a given cluster (**Figure 1H, Supplemental Figure 1I**). In the synovial lining cluster, we found the MEF2 family motifs were enriched. Mef2a, which has been shown to promote terminal differentiation of macrophages [54, 55], and it downstream targets have increased expression in this cluster. In the interstitial cluster, we found enrichment of KLF family motifs, which are well-characterized regulators of macrophage polarization [56]. While many KLF TFs are expressed in synovial macrophages, Klf3 and its downstream targets were specifically expressed in CX3CR1^-^MHCII^-^ subset. Finally, the monocyte-derived cluster exhibited significant enrichment of NFκB-p65 motif, a master regulator of innate inflammation processes through [57] and Nfkb1 emerged as a candidate TF. Taken together, we find that the four synovial macrophage populations exhibit distinct combinations of three underlying transcriptional programs.

### MHCII^+^ synovial macrophages are bone marrow derived and require CCR2 for replenishment

To further investigate the ontogeny of the synovial macrophage populations, we used the CX3CR1^ERCre/+^zsGFP fate model to assess the embryonic origins of synovial macrophages [58]. We found that almost 60% of total macrophages and almost all MHCII^-^ were GFP^+^ when mice were treated with tamoxifen at embryonic day 15, suggesting that these populations arise from erythro-myeloid progenitors in the embryo (**Figure 2A, Supplemental Figure 2A**). Next, we created parabiotic mice using B6.CD45.1 and B6.CD45.2 mice: while circulating CM are primarily self-derived, longer-lived NCM have a meaningful partner contribution (**Supplemental Figure 2B-C**). The CX3CR1^+^MHCII^+^ and CX3CR1^-^MHCII^+^ populations alone exhibited replenishment of synovial macrophages from the partner parabiotic mouse at low levels, indicating minimal turnover at steady state (**Figure 2B**), similar to a previously published study [59]. We then generated bone marrow chimeras (BMC) using B6.CD45.1 and B6.CD45.2 mice that had their ankles shielded to preserve the tissue populations. We confirmed successful reconstitution of the transplanted bone marrow as over 90% of peripheral blood cells were of donor origin (**Supplemental Figure 2D**). In this case, the MHCII^+^ populations demonstrated a significant proportion of donor-derived cells supporting a bone marrow origin (**Figure 2C, Supplemental Figure 2E**). In contrast, we observed that the vast majority (>80%) of CX3CR1^+^MHCII^-^ as well as the CX3CR1^-^MHCII^-^ macrophages were host-derived, consistent with previous reports [16, 25, 59]. These results confirm the tissue-resident and monocyte-derived origins of MHCII^-^ and MHCII^+^ populations, respectively.

**Figure 2.**
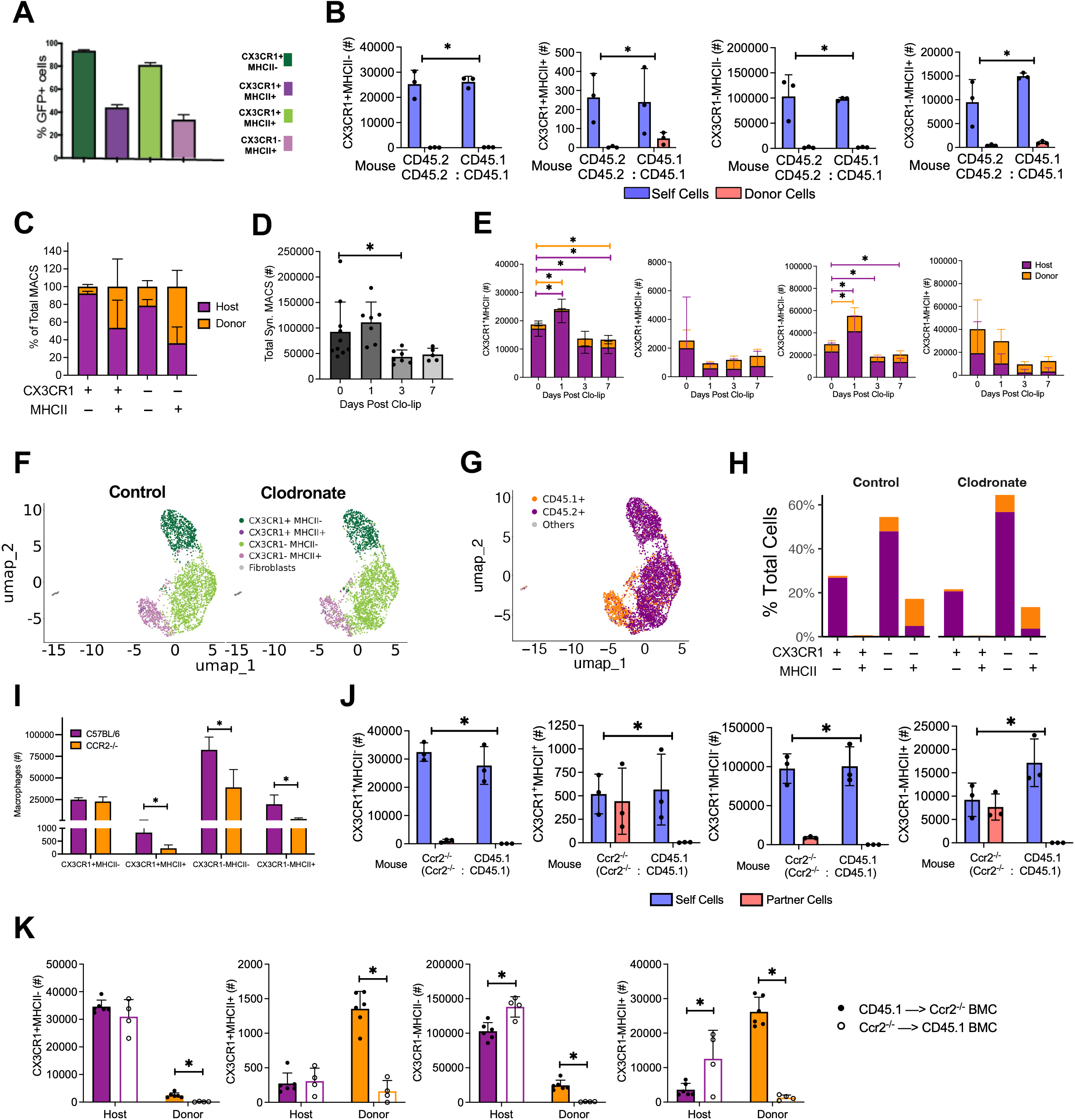
MHCII+ macrophage populations are bone marrow derived and dependent on Ccr2. **(A)** Percents of GFP+ cells of 4 macrophage subpopulations in CX3CR1^ERCre/+^zsGFP fate mapping model. **(B)** Absolute numbers of self-derived and partner-derived cells in 4 macrophages subpopulations from CD45.2:CD45.1 B6 parabiosis mice. **(C)** Percents of total macrophages of host and donor-derived cells of 4 macrophages subpopulations identified by flow cytometry in CD45.1‘*)CD45.2 bone marrow chimera mice (CD45.1‘*)CD45.2 BMC). **(D)** Absolute total numbers of macrophages identified by flow cytometry n day 0, 1, 3, 7 post clodronate-liposome (clo-lip) administration in CD45.1 à CD45.2 BMC. **(E)** Absolute numbers of 4 macrophage subsets identified by flow cytometry n day 0, 1, 3, 7 post clodronate-liposome (clo-lip) administration in CD45.1‘*)CD45.2 BMC. **(F)** UMAP depicting the distribution of cell populations in steady state (control) and day 7 post clo-lip (clodronate) CD45.1‘*)CD45.2 BMC datasets identified by CITE-seq of live CD45^+^CD11b^+^CD4^-^CD8^-^CD19^-^Ly6G^-^SiglecF^-^CD64^+^ synovial cells. **(G)** Expression of CD45.1 and CD45.2 ADT of merged steady state and clodronate CD45.1‘*)CD45.2 BMC datasets. **(H)** Percents of total cells for the 4 macrophage subsets identified by CITE-seq in the control and clodronate datasets. **(I)** Absolute numbers of 4 macrophage subsets identified by flow cytometry in B6 and CCR2^-/-^ mice. **(J)** Absolute numbers of 4 macrophage subsets identified by flow cytometry in Ccr2^-/-^:CD45.1 parabiosis mice. **(K)** Absolute numbers of 4 macrophage subpopulations identified by flow cytometry in reciprocal Ccr2^-/-^ and CD45.1 BMC mice.

We next treated the BMC mice with clodronate laden liposomes (clo-lip) and analyzed the synovial macrophage populations over time. Upon treatment, peripheral blood monocytes are severely depleted in 24 hours (**Supplemental Figure 2D**). We detected a significant reduction of total synovial macrophages at day 3 as compared to untreated (**Figure 2D**), that was primarily driven by a loss of host-derived synovial macrophages, especially in the MHCII^-^ populations (**Figure 2E**). We then performed CITE-seq on CD45^+^CD11b^+^Ly6G^-^SiglecF^-^CD64^+^ synovial macrophages from untreated (control) and day 7 following clo-lip treatment of BMC mice (**Supplemental Figure 2F**). Using label-transfer, we annotated cells based on the four populations identified in Figure 1 and validated by expression of CX3CR1 and MHCII as well as key markers (**Figure 2F and Supplemental Figure G-H**). After further annotating cells based on CD45.1/2 ADT levels, we only detect slight differences in composition or cell origin between steady-state and clo-lip macrophage populations (**Figure 2G-H, Supplemental Figure 2I**). Thus, while treatment with clodronate has a transient effect on host cells, the synovial macrophage compartment does not exhibit evidence of long-term functional changes.

Since classical monocytes require CCR2 to exit the bone marrow, CCR2^-/-^ mice lack classical monocytes in peripheral blood and tissue [60]. The numbers of CX3CR1^+^MHCII^+^, CX3CR1^-^MHCII^-^ and CX3CR1^-^MHCII^+^ synovial macrophages were significantly reduced in CCR2^-/-^ mice with depleted CM as compared to controls (**Figure 2I, Supplemental Figure 2J-K**). In CD45.1:CCR2^-/-^ parabiotic mice, CD45.1 partner cells supplemented MHCII^+^ synovial macrophages and NCM in the CCR2^-/-^ mouse although CMs stayed low (**Figure 2J, Supplemental Figure L**). The CD45.1 parabiont mouse was largely unaffected with minimal partner cell contribution. CCR2^-/-^‘*)CD45.1 BMC exhibited reduced donor derived cells in all macrophage populations. Additionally, we found that CD45.1‘*)CCR2^-/-^ BMC mice had fewer host CX3CR1^-^MHCII^-^ and CX3CR1^-^MHCII^+^ macrophages, while there were no differences in the CX3CR1^+^MHCII^-^ and CX3CR1^+^MHCII^+^ synovial macrophages as compared to CCR2^-/-^‘*)CD45.1 BMC mice, suggesting that the differentiation trajectory of CCR2-depleted macrophages in impaired (**Figure 2K, Supplemental Figure 2M**). The results indicate that MHCII+ bone-marrow derived synovial macrophages are CCR2-dependent.

### STIA is associated with an acute transcriptional response that varies across synovial macrophage populations followed by a return to near steady-state conditions

Infiltration of circulating monocytes and their subsequent differentiation into activated macrophages is a hallmark of inflammation [61]. To investigate how the four populations of synovial macrophages are altered during acute inflammatory arthritis, we utilized the STIA mouse model, which represents the effector phase of RA and requires monocytes and macrophages [62]. The clinical arthritic scores peaked at day 7 and largely returned to steady state levels by day 21 after the induction of arthritis (**Figure 3A**) consistent with previous studies [63–65]. At the same time, we observe changes in macrophage compartment composition, including a marked increase in the CX3CR1^+^MHCII^+^ populations, which reverses at day 21 (Figure 3B-C). We performed bulk RNA-seq on the sorted synovial macrophage populations at day 0, 3, 7, 13 and 21 post serum transfer to identify transcriptional changes in response to STIA. All macrophage populations were highly transcriptionally divergent at day 7 and 13, while day 21 was generally more similar compared to day 0 (**Figure S3A-B**). The CX3CR1^+^MHCII^+^ population was notable in exhibiting transcriptional changes by day 3 and maintaining changes into day 21. These data suggest that the majority of macrophages exhibit an acute transcriptional response to STIA that returns to near steady-state by resolution.

**Figure 3.**
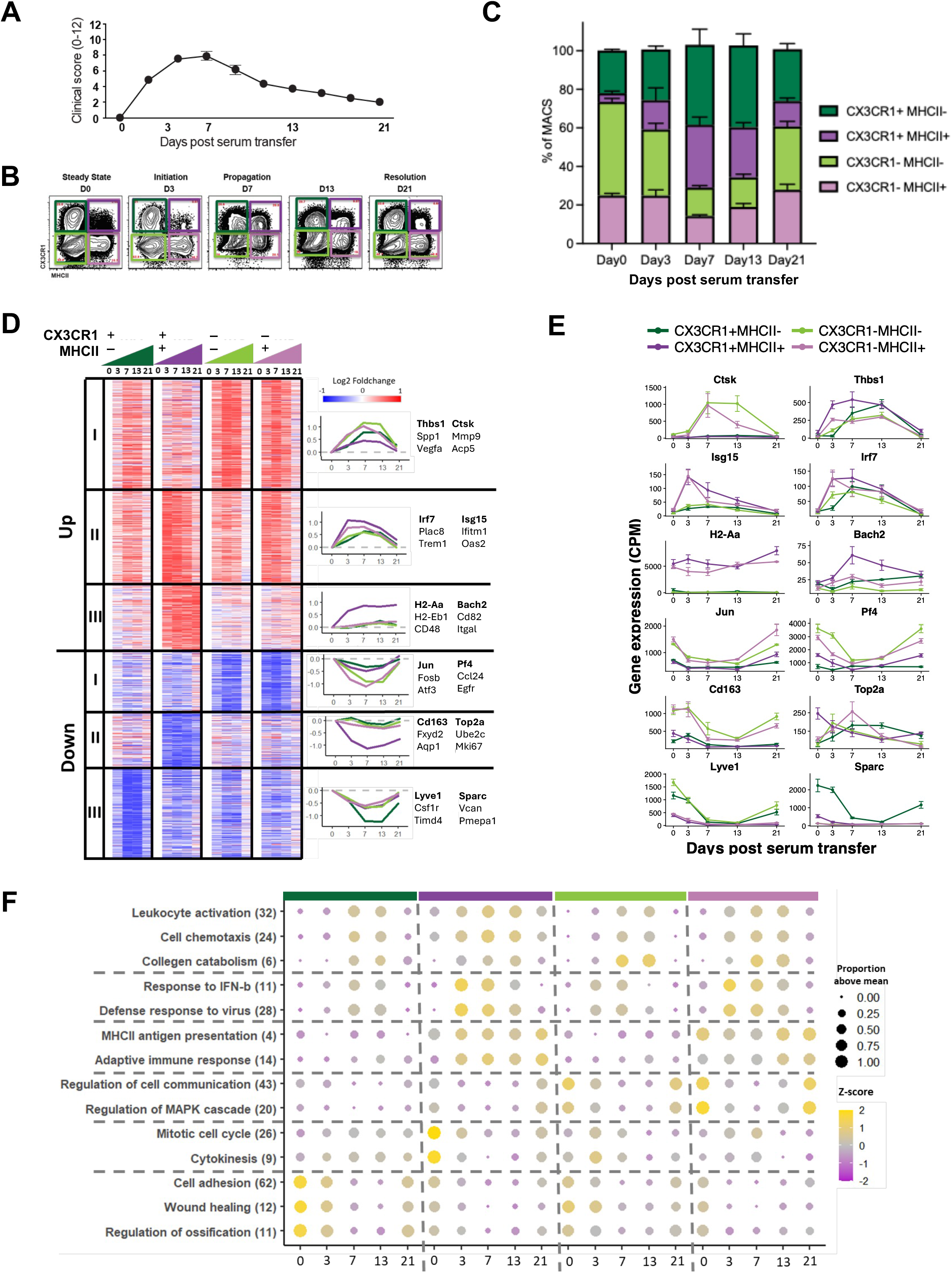
Synovial macrophage populations exhibit a mutli-dimensional response to STIA. (A) Clinical arthritic scores over the course of STIA. (B-C) Proportions of synovial macrophages subsets over time. (D) K-means clustering (K = 6) of 1772 differentially expressed genes across the STIA time course, visualized as fold-changes between the mean expressions of each timepoint and day 0 expressions. Expression trend lines are indicated on the right. (E) Gene expression of representative genes for each K-mean cluster. (F) Bubble plot showing expression trends by population/timepoint group for S select gene processes. Size of bubble indicates proportion of genes expressed at levels above their dataset average in the given group. Color scale indicates the z-score of the given group’s mean expression.

To identify temporal patterns across STIA, we defined 1772 DEGs between day 0 and any subsequent time point in at least one macrophage population. Using unsupervised K-means clustering on the relative expression compared to day 0 for each macrophage population, we defined 6 clusters of temporal gene expression patterns (**Figure 3D-F**). The first three clusters encompassed genes upregulated over the course of STIA, while the latter 3 encompassed downregulated genes. Cluster Up I included genes that are generally upregulated across populations and peaked at days 7 and 13 post-serum transfer. These included genes associated with inflammatory processes (Thbs1, Spp1, Vegfa), such as leukocyte activation and cell chemotaxis as well as those seen in osteoclasts precursors (Ctsk, Mmp9, Acp5) [19, 66, 67]. Cluster Up II consisted of upregulated genes peaking at day 3 and were the most pronounced in MHCII^+^ subsets. This cluster comprised monocyte-related genes (Irf7, Plac8, and Trem1) as well as those associated with interferon response (Isg15, Ifitm1, Oas2). Cluster Up III included genes specifically upregulated in CX3CR1^+^MHCII^+^ macrophages and persisting to 21 days after serum transfer. Genes associated with antigen presentation and adaptive immune response are overrepresented in this cluster (H2-Aa, H2-Eb1, Cd48, Bach2, CD82, Itgal).

On the other hand, Clusters Down I-III encompassed genes downregulated in response to STIA (**Figure 3D-F**). Cluster Down I consisted of genes preferentially downregulated in interstitial (CX3CR1^-^) macrophages peaking at day 7 and were enriched for signal transduction pathways, such as MAPK cascade and cAMP-mediated signaling (e.g., Atf3, Fosb, Jun) as well as genes previously observed in interstitial macrophages (Pf4, Ccl24, Egfr). Cluster Down II genes were downregulated specifically in CX3CR1^+^MHCII^+^ macrophages throughout the time course. This cluster included both genes associated with cell cycle processes (Top2a, Ube2c, Mki67) and additional interstitial macrophage genes (Cd163, Fxyd2, Aqp1). Finally, genes in Cluster Down III are primarily downregulated in the CX3CR1^+^MHCII^-^ population at day 7-13 with a rebound at day 21 and is enriched for those associated with reparative processes, such as ossification, wound healing, and cell adhesion [68–70]. This cluster also includes many genes associated with tissue-residency (Lyve1, Csf1r, Timd4) and synovial lining (Sparc, Vcan, Pmepa1).

Comparison to steady-state gene clusters confirmed that genes preferentially expressed in blood monocytes (pan-monocyte, CM, and NCM) tended to increase expression while those associated with the synovial line and interstitial phenotype tended to decrease in expression during STIA (**Supplemental Figure 3C**). Similarly, we observe an enrichment in MEF2 and KLF family motifs in the Down clusters and decreased expression of MEF2A and KLF3 targets genes which were associated with the synovial lining and interstitial clusters, respectively (**Supplemental Figure 3D-E**). Genes from the steady-state monocyte-derived cluster are observed among both the increasing and decreasing genes and NFκB, previously linked to this cluster, does not show a clear pattern. On the other hand, IRF motifs are enriched in are enriched among upregulated genes, which likely reflects an increase in monocyte contribution and/or interferon signaling. These results support the recruitment of circulating monocytes during the peak of STIA reflecting a profile distinct from steady-state monocyte derived cells and a concordant loss of tissue-resident phenotypes.

To quantify the acute versus persistent transcriptional changes in synovial macrophages, we tabulated the differentially expressed genes of day 7 and 21 relative to day 0 (**Figure S3F-G**). We found that compared to the other three subsets, CX3CR1^+^MHCII^+^ macrophages exhibit the highest number of “persistent” genes that were elevated at both day 7 and 21 post serum-transfer (Figure S3H). The number of “acute” genes, which are differential at day 7 but not day 21, is greater than those of persistent genes in both CX3CR1^-^ populations supporting a return to steady-state profiles. On the other hand, persistent genes outnumber acute, particularly in downregulation, in CX3CR1^+^MHCII^-^ macrophages suggest that post-inflammation synovial macrophages do not fully recapitulate the naïve state. In alignment with the number of genes, the slope of fold-change plots for the CX3CR1^+^MHCII^-^ and CX3CR1^+^MHCII^+^ populations was markedly higher, suggesting greater persistence, than for the other two populations.

We next utilized the CX3CR1^CreER/+^zsGFP reporting system to quantify the contribution of circulating monocytes during STIA (**Supplemental Figure 3H-I**). When treated with tamoxifen circulating (PB) monocytes from CX3CR1^CreER/+^zsGFP mice become GFP^+^ in one day but rapidly lose fluorescence due to turnover by day 7 in CM and 14 days in NCM. We first administered to steady-state CX3CR1^CreER/+^zsGFP mice on two consecutive days; all subsets exhibited subtle increases in GFP^+^ over a 21-day time course with CX3CR1^+^ populations exhibiting the highest levels overall, indicating local proliferation and minimal contribution from circulation. Next, tamoxifen was administered at Day -1 and Day 0 of the STIA time course to CX3CR1^CreER/+^zsGFP mice, which displayed large decreases in GFP positivity from day 3 to 7 suggesting significant cell replacement by circulating monocytes that would be primarily GFP^-^ by day 3. Further, we repeated the STIA time course in CX3CR1^CreER/+^zsGFP mice with tamoxifen administration at day 3 after serum-transfer. In this case, we observe an increase in GFP levels in all macrophage populations consistent with the influx of GFP^+^ circulating monocytes. Collectively, these data suggest that the acute inflammation of STIA is defined by an expansion of activated monocyte-derived macrophages followed by a resolution whereby the macrophage compartment largely returns to steady-state.

### Arthritis-associated genes are largely shared between CIA and STIA

Collagen induced arthritis (CIA), another arthritic mouse model that emulates chronic synovial inflammation rather than acute inflammation as in STIA, is potentially a better reflection of RA pathology in humans. We therefore utilized the CIA model to investigate how synovial macrophage subsets respond during chronic inflammatory arthritis [71]. In CIA, disease is established by day 41 and continues to increase in severity with no return to steady-state levels by 62 days post injection (**Figure 4A**). We sorted the four macrophage populations for bulk RNA-seq from CIA mice on day 0, 27, 41, and 62 after the first collagen injection. As in STIA, we found changes to the synovial macrophage compartment with the CX3CR1^+^MHCII^+^ population expanded starting at Day 27 and maintaining this level (**Figure 4B-C**). In contrast to STIA, the transcriptional profiles of CIA macrophage populations did not reveal an overarching trend with similar numbers of DEGs on all days (**Supplemental Figure 4A-B**). Since day 41 CIA demonstrated the highest correlation with peak inflammation at day 7 STIA, we focused out analyses at this timepoint (**Supplemental Figure 4C**). We found a high overlap between DEGs by population in day 7 STIA and day 41 CIA (**Figure 4D**). Moreover, CIA DEGs corresponded to STIA clusters from Figure 3 in a population-specific manner and these genes exhibited similar expression patterns across model but on a lengthened timeline in CIA (**Figure E, Supplemental Figure 4D**). Similarly, previously described “acute” genes in STIA were annotated as chronic (differential in both day 41 and day 62) along with the majority of CIA DEGs (**Supplemental Figure 4E-F**). This is reflected in the slope increase of fold-changes across all macrophage populations in CIA compared with STIA (CX3CR1^+^MHCII^-^: 0.36 to 0.57; CX3CR1^+^MHCII^+^: 0.44 to 0.61; CX3CR1^-^MHCII^-^: 0.19 to 0.48; CX3CR1^-^MHCII^+^: 0.19 to 0.46). Overall, we observe that the transcriptional profile of macrophages in CIA is consistent with a lengthened inflammatory response akin to STIA.

**Figure 4.**
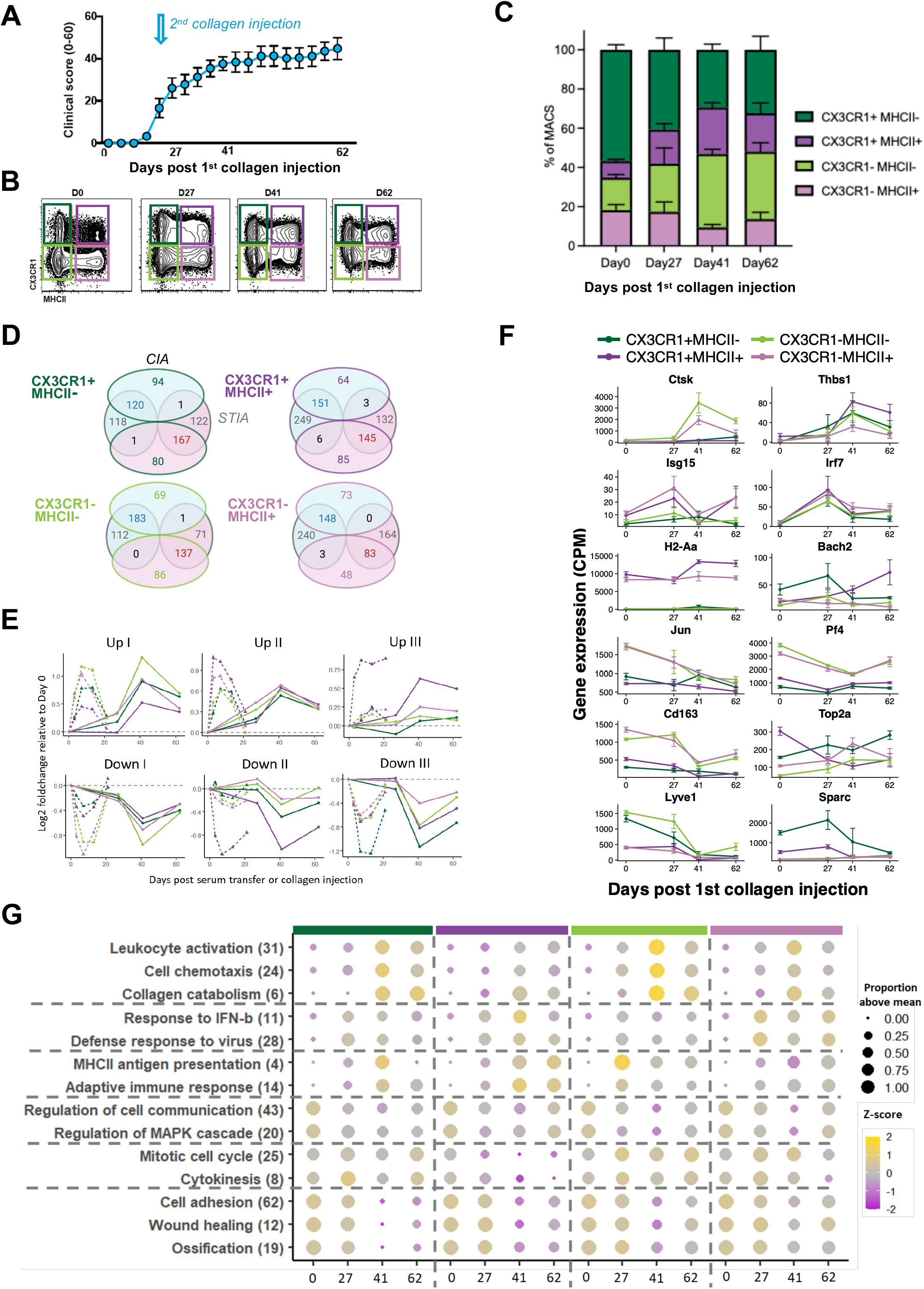
Macrophage response in CIA shares high overlap with STIA. (A) Clinical arthritic scores over the course of CIA. (B-C) Proportions of synovial macrophages subsets over time. (D) Average log expression fold changes relative to Day 0 for STIA cluster genes in macrophage populations from STIA and CIA models. (E) Venn diagrams showing overlap in up-regulated (blue) and down-regulated (red) DEGs for each macrophage population in CIA (colored by populations) vs. STIA (grey). (F) Gene expression of representative genes that are DEGs in both CIA and STIA (G) Bubble plot showing expression trends by population/timepoint group for select gene processes. Size of bubble indicates proportion of genes expressed at levels above their dataset average in the given group. Color scale indicates the z-score of the given group’s mean expression.

In addition to general patterns, we find high overlap in the specific genes and pathways that are regulated in response to CIA and STIA (Figure 4F). Specific genes that are generally upregulated in both CIA and STIA include those involved in cell chemotaxis (e.g., Spp1, Lgals3), leukocyte activation (Thbs1, Ccr2), and collagen catabolism (e.g., Ctsk, Mmp9) (Figure 4G). The upregulation of Type I interferon genes, a major characteristic of CX3CR1^+^MHCII^+^ subset during STIA time course, is more subtle CIA, possibly due to the longer timeline missing an early peak. However, we did observe increased expression of monocyte-related (Irf7, Plac8), antigen presentation (H2-Aa, H2-Eb1), and adaptive immune response (Bach2) genes in at least one macrophage population. CIA also conserved the downregulation of genes associated with cell communication across macrophage populations (Jun, Atf3, Fosb) and cell division in CX3CR1^+^MHCII^+^ macrophages (Top2a, Ube2c, Mki67). There was a clear decrease in expression of homeostatic and reparative processes including cell adhesion, wound healing, and ossification (Cd9, Serpine 1, Vcan, Bmp2, Gas6, Egr2). Collectively, we observe a strong conservation in subset-specific expression of biological pathways regardless of arthritis model.

Furthermore, CIA exhibits analogous trends as STIA with respect to steady-state macrophage population signatures. Genes generally associated with tissue-residency (Lyve1, Timd4) and specifically with synovial lining (Sparc, Pmepa1) or interstitial (Pf4, Cd163) phenotypes were decreased in expression while those associated with monocytes (Irf7, Plac8, Ccr2, Cd52) tended to be increased. Similarly, we observe a downregulation of downstream target genes for TFs, MEF2A and KLF3 (**Supplemental Figure S4F**), which were previously attributed to maintenance of steady state synovial macrophage functions. Therefore, despite differences in the timeline and etiology, synovial macrophages exhibit transcriptionally similar responses to CIA and STIA.

### STIA gives rise to novel monocyte-derived macrophage subpopulations

In order to further investigate heterogeneity and ontogeny of macrophages in inflammatory arthritis, we analyzed synovial macrophages from B6.CD45.1‘*)B6.CD45.2 BMC at days 0, 1, 7, 14, and 56 post-STIA (**Figure 5A, Supplemental Figure 5A**). By flow cytometry, we observed a dramatic increase in donor-derived cell numbers across all populations by the plateau phase of inflammation. Using CITE-seq, we defined 8 *de novo* gene expression clusters spanning across time points as well as applying label transfer to annotate cells with the steady-state identities and using CD45.1/CD45.2 ADT expression to annotate host vs. donor origins (**Figure 5B-C, Supplemental Figure 5B-H**). We found that the clusters 0 and 1 align with steady-state populations CX3CR1^-^MHCII^-^ and CX3CR1^+^MHCII^+^ respectively, exhibit CD45.2 expression indicating they are host-derived, and are depleted during the inflammatory phase of STIA (day 7-14). Clusters 4 and 7 are also most abundant in steady-state and are host-derived: their profiles seem to match the Clec10a and Arg1-expressing subpopulations, respectively, from Figure 1. Clusters 3 and 5 are present across the STIA time course though their shift in location within the UMAP suggests underlying transcriptional changes. Moreover, cluster 3 is donor-derived and expresses MHCII-related genes although it is first annotated as CX3CR1^-^MHCII^+^ in steady state but switches to CX3CR1^+^MHCII^+^ on day 7 and back again on day 56. Cluster 5 is host-derived at steady-state, but donor-derived in inflammation: continuous expression of Ctsk combined with induced expression Acp5, Ocstamp and Mmp9 suggests these are osteoclast precursors observed previously in single-cell RNA-seq analyses of murine synovial macrophages[19, 24]. Finally, clusters 2 and 6 are both unique to the inflamed joint. Cluster 2 is donor-derived and expresses activation-associated genes such as Spp1, Thbs1, Vegfa, and Mmp14. On the other hand, cluster 6 is host-derived and expresses Clec10a, possibly reflecting a STIA-responsive version of interstitial macrophages. Interestingly at Day 56 when inflammation is resolved we largely observe a reappearance of steady-state populations though they exhibit a mix of donor and host origins.

**Figure 5.**
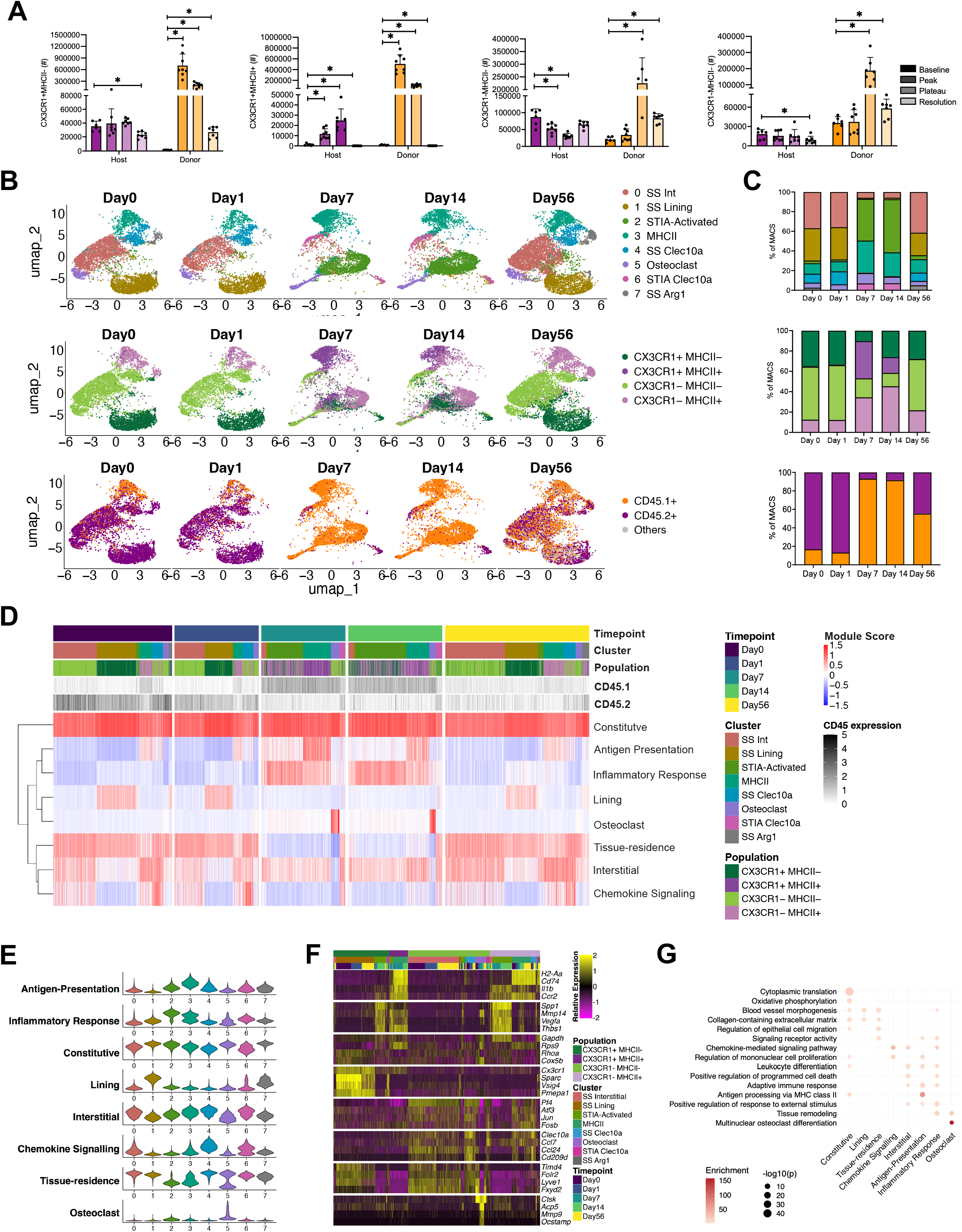
Acute arthritis is defined by an influx of monocyte-derived macrophages and a loss of distinct macrophage phenotype. **(A)** Absolute numbers of host- and donor-derived cells macrophage subsets in baseline, peak, plateau, and resolution stages of K/BxN serum transfer induced arthritis (STIA) CD45.1àCD45.2 BMC mice. **(B)** UMAP depicting 8 clusters, 4 macrophage subpopulations, and CD45 annotation in Day0, day1, day7, day14, day56 STIA CD45.1àCD45.2 BMC identified by CITE-seq of liveCD45^+^CD11b^+^CD4^-^CD8^-^CD19^-^Ly6G^-^SiglecF^-^CD64^+^ synovial cells. **(C)** Compositions of 8 clusters, 4 macrophage subpopulations, and CD45.1+/CD45.2+ cells in Day0, day1, day7, day14, day56 STIA CD45.1àCD45.2 BMC CITE-seq datasets. **(D)** Module score heatmap of the 8 identified gene modules for cluster stratified by timepoint and population. **(E)** Module gene expression of 8 clusters. **(F)** Relative gene expression heatmap of selected markers of 8 module genes for population stratified by cluster and timepoint. **(G)** GSEA pathway enrichments result of 8 module genes.

To better understand how transcriptional profiles of synovial macrophages may change within populations across STIA, we performed a modular analysis to identify co-regulated genes (**Figure 5D-G, Supplementary Figure 5I)**. We identified 8 modules and scored cells based on their expression. The Constitutive module includes genes that are ubiquitously expressed across synovial macrophage populations, origin, and inflammatory state (Gadph, Rps9, Rhoa, and Cox5b), such as those involved in ongoing processes like translation and oxidative phosphorylation. The Antigen Presentation module expressed MHCII and adaptive immune response genes and is exclusively expressed in MHCII^+^ annotated populations (H2-Aa, Cd74, Il1b, Cbr2). The Inflammatory Response module is primarily expressed by clusters 2 and 3 in inflammation and includes genes previously associated with activation in our arthritis models (Spp1, Mmp14, Vegfa, Thbs1). The Lining (Cx3cr1, Sparc, Vsig4, Pmepa1) and Tissue-residence (Timd4, Folr2, Lyve1, Fxyd2) modules both seem to disappear during inflammation and reappear in resolution. In contrast, the Interstitial module (Pf4, Atf3, Jun, Fosb) persists in the inflammatory phase and indeed expands in the inflammatory phase. Interestingly, the Chemokine Signaling module 7 (Clec10a, Ccl7, Ccl24, Cd209) is largely opposite to the Inflammatory Response module with broad expression across steady-state but only host-derived cells in inflammation. Lastly, the Osteoclast module contains osteoclast differentiation genes Ctsk, Acp5, Mmp9, and Ocstamp but is only observed in the midst of inflammation. These transcriptional modules demonstrate a high concordance with the gene expression patterns identified from bulk RNA-seq data (**Supplemental Figure 5J-K**). Taken together, these results indicate that synovial macrophage exhibit novel transcriptional signatures between steady-state and inflammation.

### Transcriptional signatures of arthritis are conserved at the cellular level in the KRNAg7 spontaneous model

The KRNAg7 model of inflammatory arthritis (KRN) is distinct from CIA and STIA as synovitis develops spontaneously at around 3-4 weeks of age. Thus, synovial macrophages in this model likely undergo long-term remodeling. To determine how macrophage heterogeneity in KRN compares with our other models, we performed single-cell RNA-seq on CD64+ synovial cells (**Supplemental Figure 6A)**. We found a higher level of neutrophil contamination than seen previously. *De novo* clustering defined 8 clusters of which 4 are clearly identified as macrophages: cluster 0 (Trem2, Fxyd2, Pf4) comprises cells annotated as both tissue-resident synovial lining (CX3CR1^+^MHCII^-^) and interstitial (CX3CR1^-^MHCII^-^) cells, clusters 1 and 4 (Ccr2, Ly6c2, Plac8) annotated as infiltrating macrophages (CX3CR1^+^MHCII^+^), and cluster 2 (H2-Aa, H2-Eb1, Il1b) annotated as monocyte-derived (CX3CR1^-^MHCII^+^) (**Figure 6A, Supplemental Figure 6B-C)**. Label transfer of steady-state populations annotations confirms these relationships (**Figure 6B-C, Supplemental Figure 6D**).

**Figure 6.**
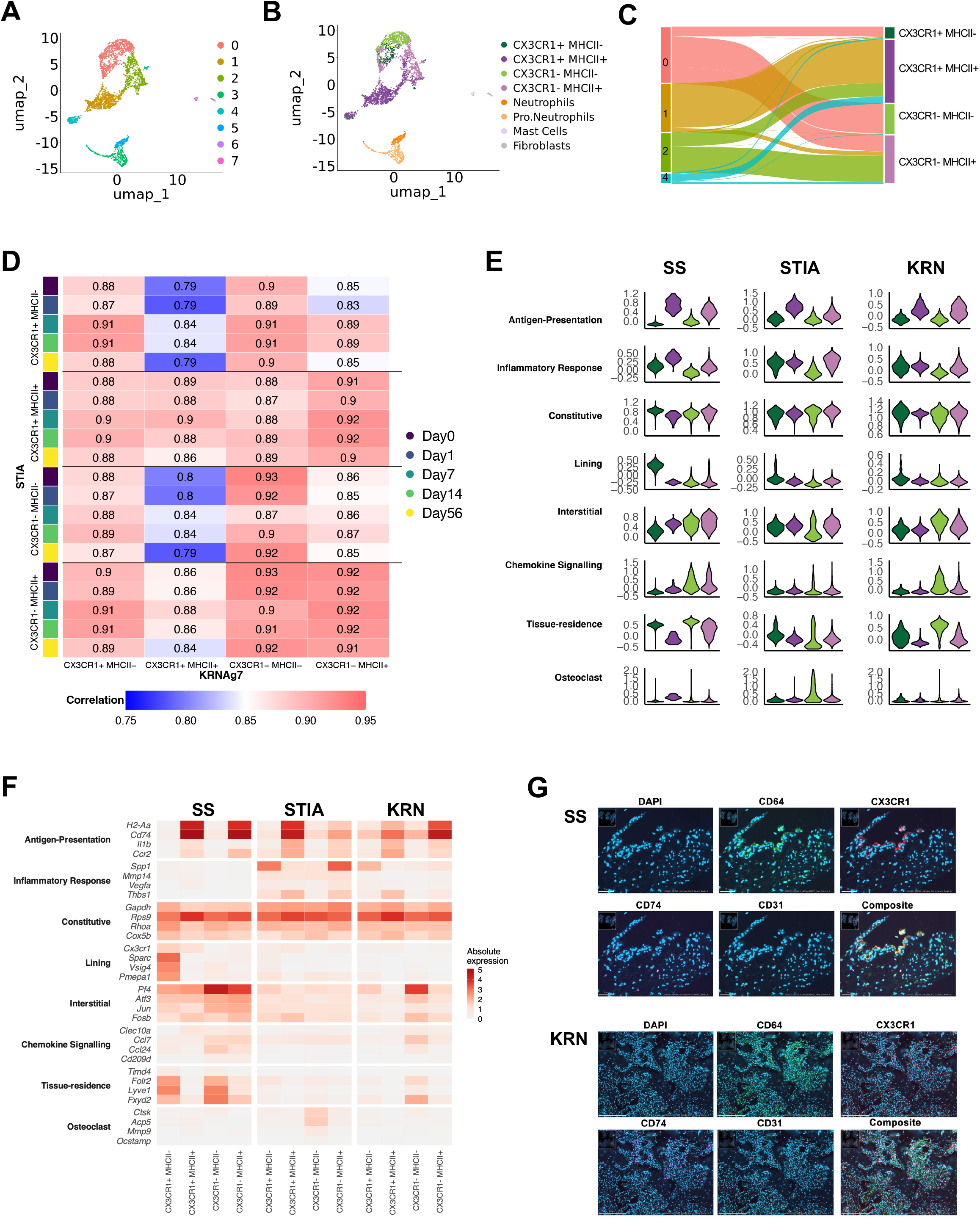
Macrophages from KRNAg7 exhibit inflammatory signatures in tandem with interstitial remodeling. **(A)** UMAP depicting 8 clusters identified in KRNAg7 mice by CITE-seq of liveCD45^+^CD11b^+^CD4^-^CD8^-^CD19^-^Ly6G^-^SiglecF^-^CD64^+^ synovial cells. **(B)** Annotation of cell types. **(C)** Sankey plot of the contribution of 4 macrophage subpopulation from clusters in macrophage subset dataset. **(D)** Correlation between 4 macrophage subpopulations from KRNAg7 CITE-seq data and STIA CD45.1‘*)CD45.2 BMC datasets stratified by timepoint. **(E)** Expression of module scores from the 8 module gene across 4 macrophage subpopulations in steady state B6 (SS), Day 7 STIA BMC (STIA), and KRNAg7 (KRN) datasets. **(F)** Expression of representative genes from the 8 module genes across 4 macrophage subpopulations in SS, STIA, and KRN datasets. **(G)** Immunofluorescence staining of CD64, CX3CR1, CD74, CD31 in SS and KRN synovial tissue.

KRN macrophages express a mixture of genes we previously associated with steady-state and inflammation. While the KRN CX3CR1^+^MHCII^-^ population most closely resembles its counterpart in the inflammatory phase of STIA, CX3CR1^-^MHCII^-^ exhibit a higher correlation with their steady-state/resolution counterpart (**Figure 6D).** Similarly, the expression of the transcriptional modules identified in STIA is largely consistent with that seen on day 7 (**Figure 6E-F, Supplemental Figure 6E)**. The Antigen Presentation and Inflammatory response exhibit consistent patterns in MHCII^+^ populations. The Constitutive module is ubiquitous in KRNAg7. The Lining module is virtually absent. On the other hand, the Tissue-residence, Interstitial, and Chemokine Signaling modules is more highly expressed in the CX3CR1^-^MHCII^-^ population; this suggests that more of the host-derived phenotype is preserved in the long-term inflammation of KRNAg7. Interestingly, we see very little expression of the Osteoclast module. The KRN model also agrees with the population-specific expression patterns identified in bulk data (**Supplemental Figure 6F**). Consistent with the transcriptional signatures, we observe a loss of CX3CR1+ macrophages and gains of CD74+ (MHCII+) macrophages in KRN vs. steady-state synovial tissue. Taken together, these results indicate that similar inflammatory processes are active in KRNAg7 joints though the tissue-resident phenotype is partially maintained.

## Discussion

In this study, we compared synovial macrophage ontogeny and transcriptional heterogeneity across steady-state and various models of inflammation. We propose that surface level expression of CX3CR1 and MHCII captures the major macrophage populations. These populations generally remain distinct in inflammation but share expression of activation pathways that are conserved across arthritis models. The donor-derived macrophage proportion is expanded in the inflamed joint but these cells are capable of adapting steady-state profiles with resolution. Our results provide the first comprehensive characterization of the synovial macrophage compartment across 3 common inflammatory arthritis models.

We focus on 4 major populations of synovial macrophages: synovial lining (CX3CR1^+^MHCII^-^), tissue-resident interstitial (CX3CR1^-^MHCII^-^), monocyte-derived (CX3CR1^-^MHCII^+^), and infiltrating (CX3CR1^+^MHCII^+^). Our results support those of Culemann et al demonstrating that CX3CR1^+^ are long-lived cells with a distinct transcriptional profile[19]. As we published previously [16], MHCII marks monocyte-derived cells at steady-state with the CX3CR1^+^MHCII^+^ demonstrating the largest turnover. In addition, we show that these populations are Ccr2-dependent at steady-state. As part of our analysis we define smaller subpopulations, such as Clec10a and Aqp1-expressing interstitial cells. However, these are less robust across models and are not always recapitulated in other single-cell studies [19, 23, 24, 47]. In our own data, specific Aqp1 cells were not identified in STIA and Clec10a-expressing cells were found across clusters. In addition, our study does not include CD64- myeloid cells in the joint, such as the tissue-resident monocyte-lineage cell (TRMC) population we have previously described [25, 26]. Finally, we characterized other surface markers that vary between populations including CD163, CD11a, CD11c, and CD106 that may be used in the future to isolate these populations.

Our study represents the most in-depth investigation yet of the synovial macrophage compartment in inflammatory arthritis. We found activation signatures that were conserved across three different models. Moreover, we observed an upregulation of monocyte-related pathways with inflammation and a decrease in tissue-residency, particularly for the synovial lignin phenotype. Using bone marrow chimerism and single-cell approaches, we confirmed that there is an influx of bone marrow derived cells in peak inflammation that persists into resolution. Moreover, we identified novel arthritis-associated subpopulations and transcriptional modules within previously annotated populations. In future models, a combination of population annotation and transcriptional module expression can be used to uniquely identify macrophage state.

It is difficult to apply our results to clinical samples from RA patients. Neither CX3CR1 nor MHCII expression in mice matches humans. However, we do observe some alignment between in gene expression across the tissue-residency (MHCII-) vs. monocyte-derived (MHCII^+^) axis. MERTK, FOLR2, and CD206 are likely to delineate long-lived, tissue-resident macrophages. Among these cells, both TREM2 and NUPR1 have been connected with synovial lining macrophages and are decreased in rheumatoid arthritis patients [13, 72]. Other tissue-resident cells in rheumatoid arthritis express LYVE1 and may therefore represent long-lived interstitial cells. On the other hand, CD48 has been linked to the more monocyte-derived phenotype of synovial macrophages that can be found across conditions. A subset of these are high in S100A8, S100a9, and S100a12 and appear to be monocyte-like cells that are increased in RA [13, 72]. Categorizing human macrophage by pathway rather than specific genes will facilitate a deeper understanding of their role in RA.

Our analysis indicates conflicting roles for Arg1 in synovial macrophages across inflammatory time courses. Initially, we observe an CX3CR1^-^MHCII^+^ Arg1-expressing subpopulation at steady-state co-expressed with Flt1 and Anxa8. However, we observe and increase in Arg1 expression in STIA that is co-expressed in synovial macrophages at peak inflammation with Spp1, Thbs1, and Vegfa. With resolution, this arthritis-associated subpopulation disappears while the steady-state subpopulation reappears, but only in host-derived cells. These results add complexity to the canonical idea that Arg1 is associated with a reparative phenotype. In addition, our prior work demonstrates opposing trends for Arg1-expressing macrophages between sexes in aging [24]. Further study is needed to elucidate the role of the Arg1-expressing arthritis-associated subpopulations in disease.

Osteoclast precursor cells represent another unique subpopulation among synovial macrophages. As seen previously, we observe Ctsk-expressing cells in the CX3CR1-MHCII-population at steady-state (Dapas et al, Culemann et al). Interestingly, these cells only obtain the full profile of osteoclast precursors with expression of Mmp9 and Ocstamp in inflammation. We have also previously observed similar cells in the CD64^-^ compartment of the synovial myeloid cells and other groups have suggested that these osteoclast precursors play a critical role in arthritis [26, 66]. A better understanding of how these cells change in inflammation may shed light on bone erosion in arthritis.

The genomic and immunological approaches used in this study present some inherent limitations. First, we are unable to track specific cells across time courses and can therefore only infer relationships based on transcriptional similarity. Similarly, both ADT and flow-based assessment of surface marker expression are imperfect measures and introduce error in population definitions. Thus, there are limits to the precision of experiments and how well the results align across experiments. Finally, the synovial macrophage compartment exhibits variability across individual mice, especially in models of inflammation. We mediate this limitation to the best of our ability by using replicates for flow and pooling across animals for single-cell approaches.

Standardizing the classification of synovial macrophages in mouse models of inflammatory arthritis will enable the field to better interpret results across studies. By identifying conserved activation pathways, our studies overcome the confounding variables of individual models and support a common mechanism of synovitis that is likely to reflect human disease. Moreover, we highlight specific macrophage subpopulations and transcriptional modules as potential therapeutic targets. With myeloid-specific reporter mice and cre-flox models, researchers can further dissect the functional implications for pathogenesis [73]. Future work is needed to identify the transcription factors regulating the activity of macrophages in inflammatory arthritis. With this knowledge, we can develop models to specifically modulate macrophages and enable translation into the treatment of rheumatoid arthritis.

---

**Supplemental Figure 1.**
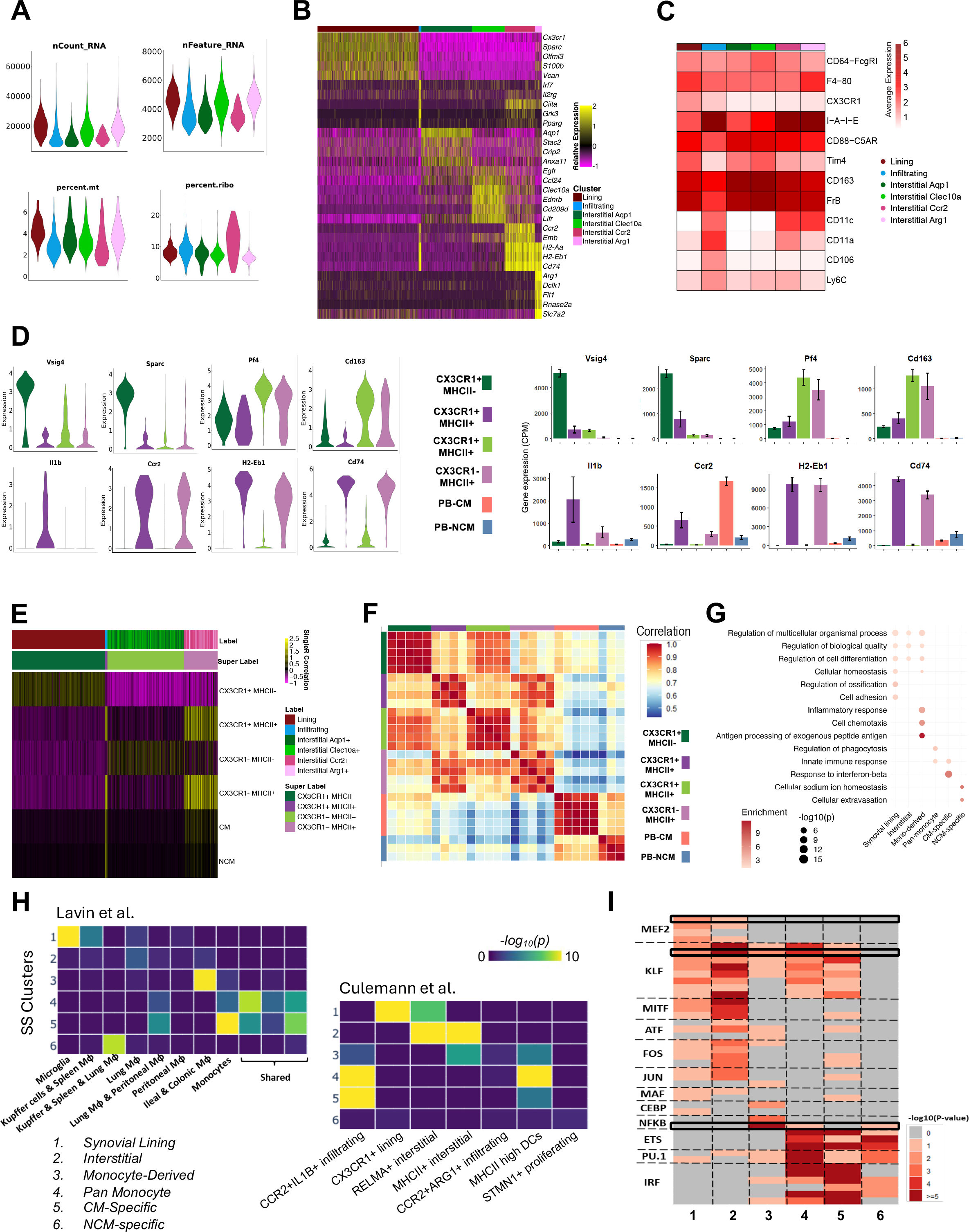
**(A)** CITE-seq quality control (QC) results of nCount_RNA, nFeature_RNA, percent.mt, and percent.ribo of 6 macrophage subpopulations in B6 CITE-seq dataset. **(B)** Heatmap of top 5 differentially expressed genes (DEG) of 6 macrophage subpopulations. **(C)** Heatmap of selected ADT markers in 6 macrophage subpopulations. **(D)** Violin plot of the key genes expression in CITE-seq identified 4 macrophage subpopulations and bulk RNA-seq analyzed 4 macrophages subpopulation as well as PB CM and NCM. **(E)** Correlation of CITE-seq identified 4 macrophage subpopulations based on CX3CR1 and MHCII expression with annotated macrophages subpopulations based on bulk RNA-seq gene expression profiles. **(F)** Correlation of gene expression across each bulk RNA-seq samples. **(G)** Gene set enrichment analysis (GSEA) results of pathway enrichments in macrophage subsets and PB monocytes. **(H)** Enrichment significance of key genes identified from 6 macrophage subpopulations in published multi-tissue macrophage datasets (Lavin et al. (REF)) and synovial macrophage subsets (Culemann et al. (REF)). **(I)** Heatmap displaying the identified differential transcriptional factors (TF) in 6 macrophage subpopulations.

**Supplemental Figure 2.**
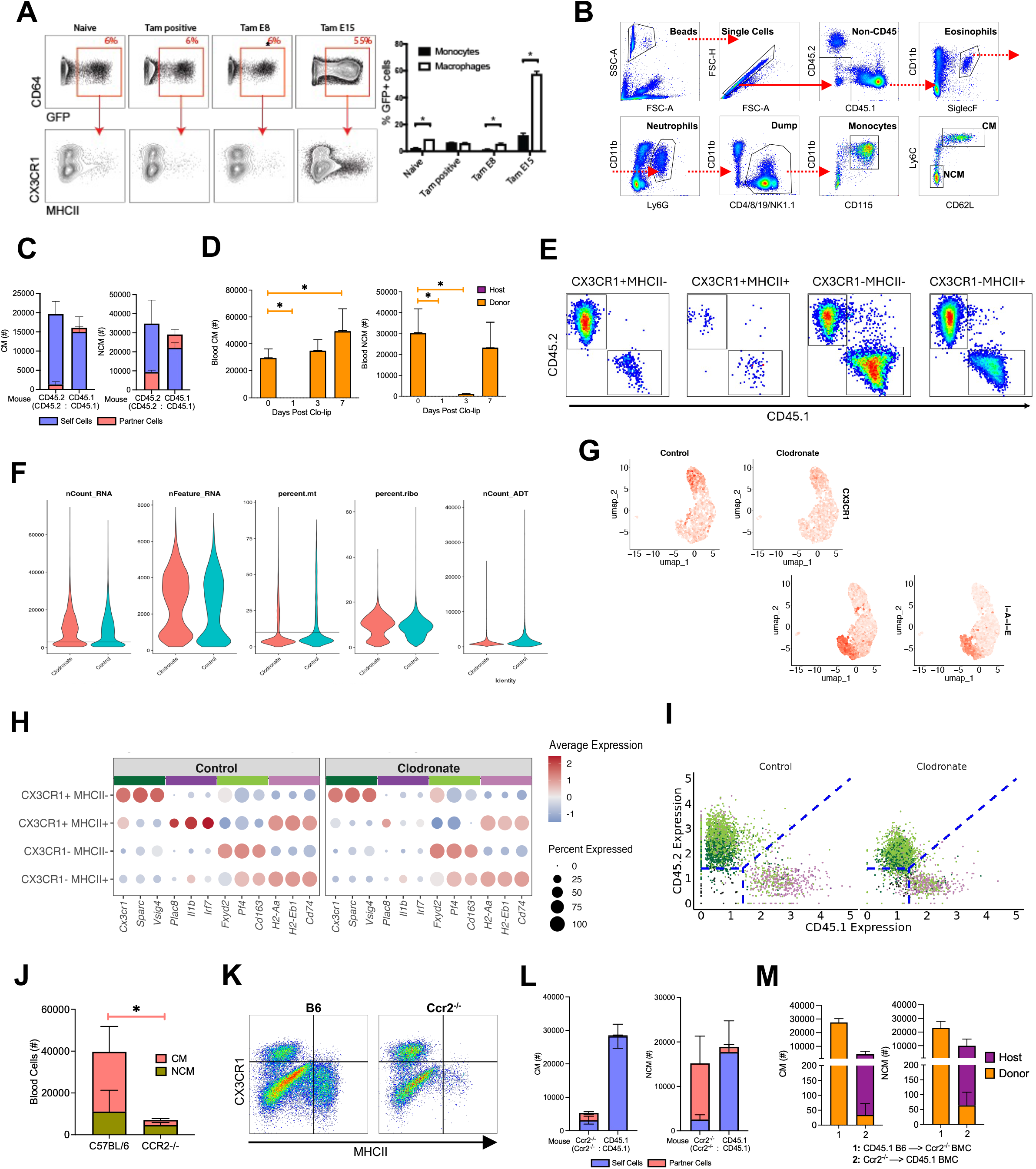
**(A)** CX3CR1 and MHCII expression in GFP+ cells, and percents of GFP+ cells in monocytes and macrophages from CX3CR1^ERCre/+^zsGFP fate mapping mice treated with tamoxifen during adulthood and embryonic stage. **(B)** PB flow cytometry gating plot. **(C)** Absolute numbers of CM and NCM in CD45.2:CD45.1 parabiosis mice. **(D)** Flow cytometry example of CD45.1 and CD45.2 expression of 4 macrophage subpopulations in Day 0 CD45.1 à CD45.2 BMC. **(E)** Absolute numbers of PB CM and NCM in CD45.1 à CD45.2 BMC mice following clo-lip administration. **(F)** CITE-seq QC results of nCount_RNA, nFeature_RNA, percent.mt, percent.ribo, nCount_ADT in day 0 (control) and day 7 post clo-lip (clodronate) CITE-seq datasets. **(G)** Feature plots of antibody derived tag (ADT) CX3CR1 and I-A-I-E (MHCII) Expression. **(H)** Average gene expression of selected gene markers as in Figure 1B of 4 macrophage subpopulations in control and clodronate CITE-seq datasets. **(I)** The thresholds of identifying CD45.1+ and CD45.2+ cells in control and clodronate datasets. **(J)** Absolute blood numbers of CM and NCM in B6 and Ccr2^-/-^ mice. **(K)** Flow cytometry example of 4 macrophage subsets in B6 and Ccr2^-/-^ mice. **(L)** Absolute numbers of CM and NCM in CD45.2:CD45.1 parabiosis mice. **(M)** Absolute numbers of CM and NCM in CD45.1 B6 and Ccr2^-/-^ reciprocal BMC mice.

**Supplemental Figure 3.**
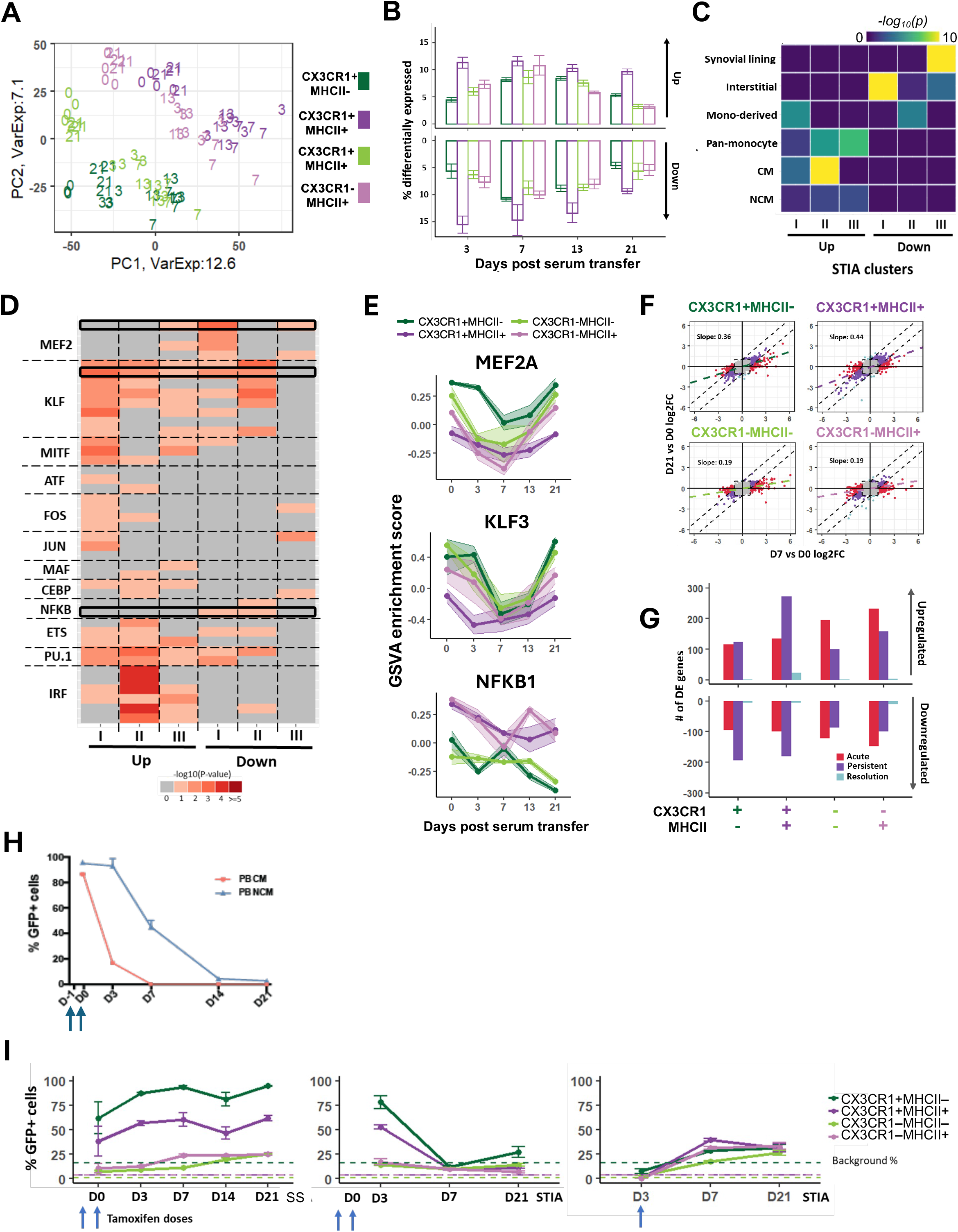
(A) PCA of gene expression profiles between replicates from STIA time course. Text indicates number of days after serum transfer. (B) Quantification of up- and down-regulated genes between individual time points of STIA. (C) Fraction of gene overlap between steady state and STIA K-means clusters, with sizes of STIA clusters as denominators. (D) Significance of DNA binding motif enrichments for select TF families across STIA k-means clusters. The color scale represents -log10(p-value) as computed by HOMER. (E) GSVA-inferred expression scores for downstream target genes of MEF2A, KLF3, and NFKB-p65 from Dorothea database across macrophage subsets and STIA time course. (F) Scatter plots of expression fold changes relative to Day 0 STIA for Day 7 and Day 21. (G) Quantification of acute (Day 7 only; red), persistent (Days 7 & 21; purple), and resolution (Day 21 only; light blue). (H) % of GFP+ cells across peripheral blood (PB) monocyte populations following tamoxifen administration to CX3CR1^CreER/+^zsGFP mice. CM = Classical Monocytes, NCM= Non-classical Monocytes (I) % of GFP+ cells across macrophage subsets following tamoxifen administration to CX3CR1^CreER/+^zsGFP mice: Steady-state mice were administered tamoxifen doses at Day 0 and Day -1 relative to first time point (left), STIA mice were administered tamoxifen doses at Day 0 and Day -1 (middle), or Day 3 relative to serum transfer. The background % of GFP expression in non-tamoxifen treated control CX3CR1^CreER/+^zsGFP mice are indicated in dashed lines.

**Supplemental Figure 4.**
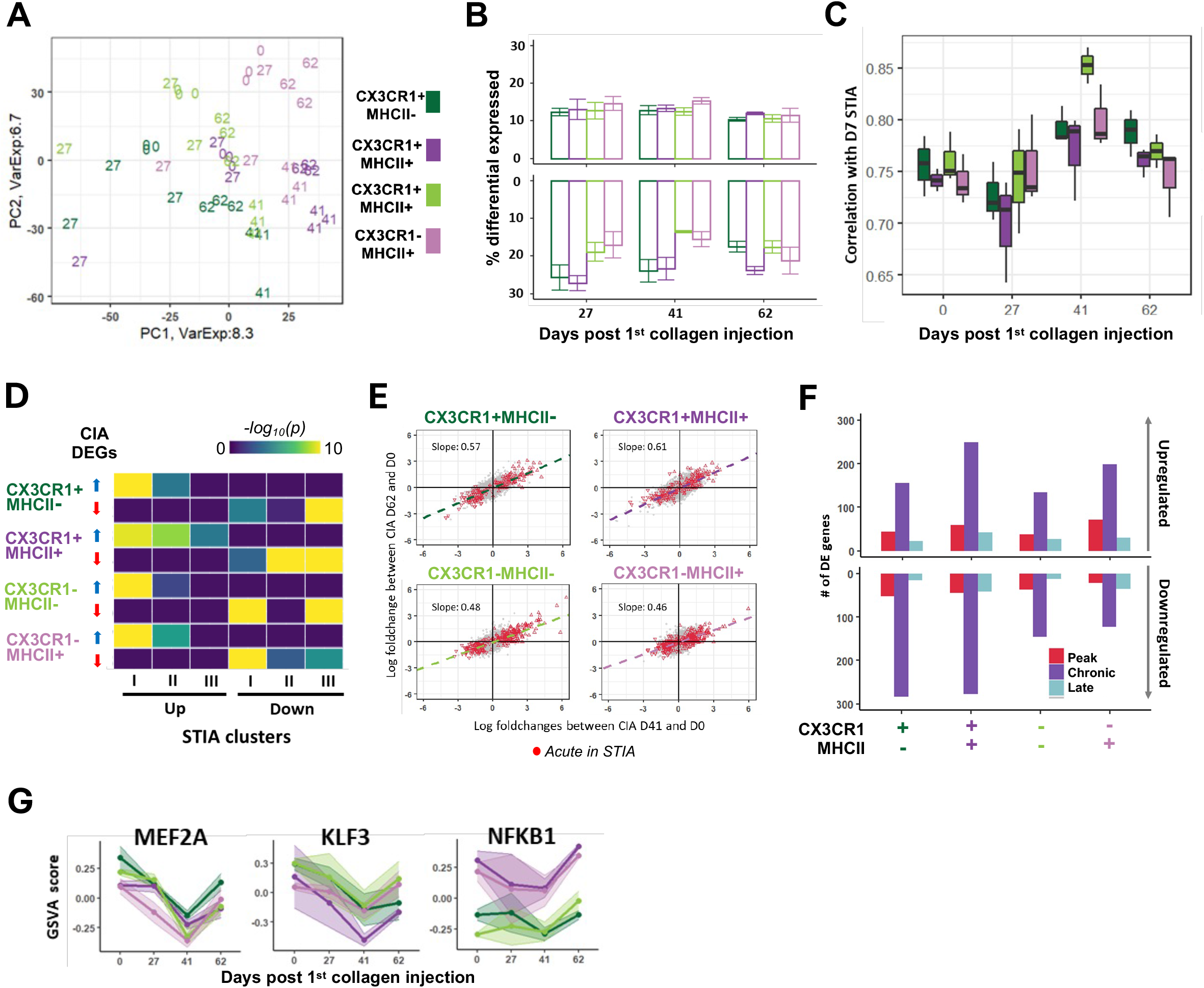
**(A)** PCA of gene expression profiles between replicates from CIA time course. Text indicates number of days after serum transfer. (B) Quantification of up- and down-regulated genes between individual time points of STIA. (C) Correlations of gene expression profiles between day 7 STIA subpopulations and CIA populations over time. (E) Scatter plots of CIA Day 41 and Day 62 expression fold changes relative to Day 0. Red dots represent STIA acute genes defined in Figure S3. (F) Quantification of peak (Day 41 only; red), chronic (Days 41 & 62; purple), and resolution (Day 62 only; light blue) genes. (G) GSVA-inferred expression scores for downstream target genes of MEF2A, KLF3, and NFKB-p65 from Dorothea database across macrophage subsets and STIA time course.

**Supplemental Figure 5.**
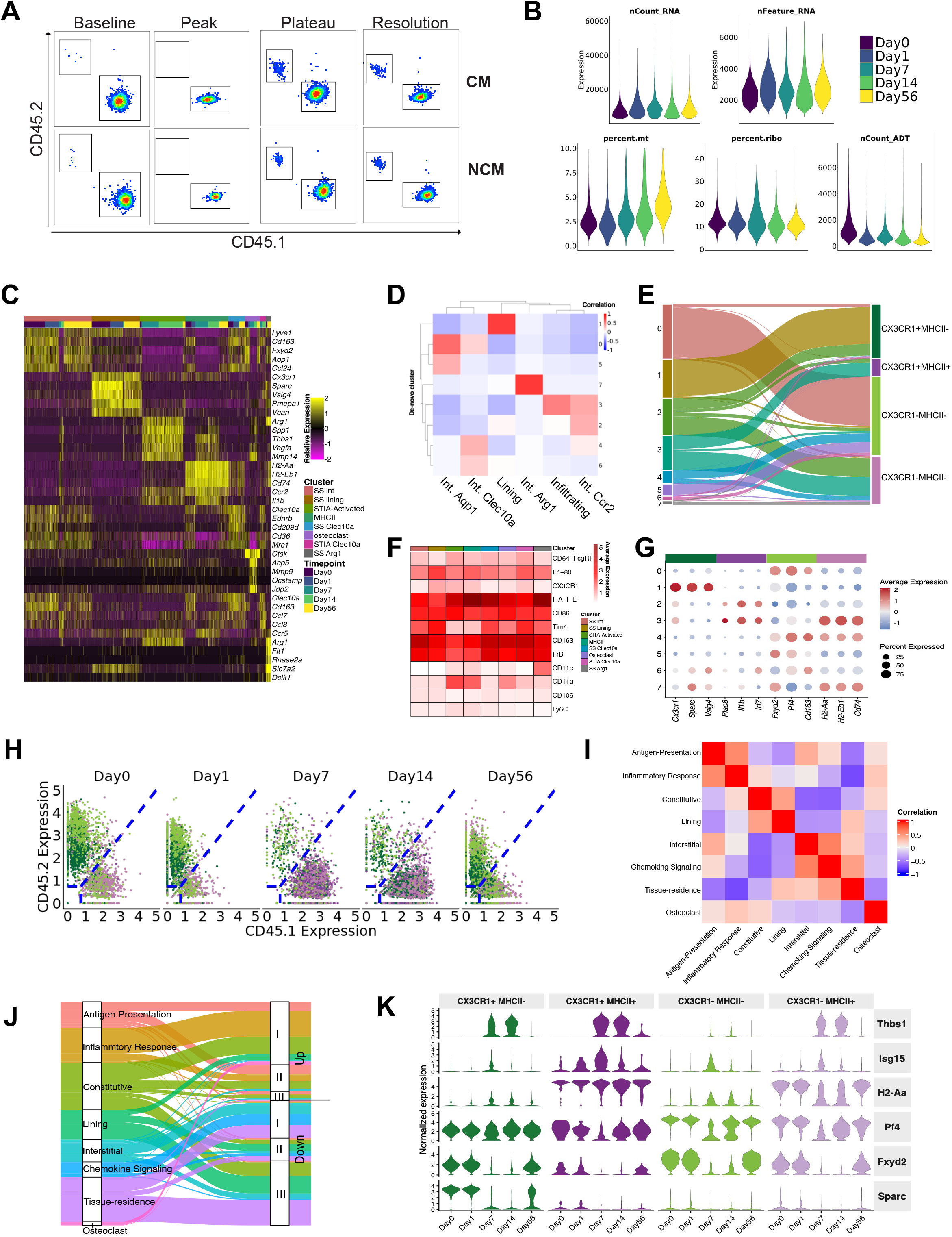
**(A)** Flow cytometry example of CD45.1 and CD45.2 expression for CM and NCM in baseline, peak, plateau, and resolution stages of STIA CD45.1àCD45.2 BMC mice. **(B)** CITE-seq QC results of nCount_RNA, nFeature_RNA, percent.mt, percent.ribo, nCount_ADT in day0, day1, day7, day14, day56 STIA CD45.1àCD45.2 BMC mice CITE-seq datasets. **(C)** Heatmap of selected gene markers for each cluster stratified by timepoint. **(D)** Heatmap of phi correlation matrix between cluster and label transfer annotation based on Figure 1A in merged day0, day1, day7, day14, day56 STIA CD45.1àCD45.2 BMC datasets. **(E)** Sankey plot of the contribution of population from cluster in merged day0, day1, day7, day14, day56 STIA CD45.1àCD45.2 BMC datasets. **(F)** ADT heatmap of cluster stratified by timepoint. **(G)** Bubble plot gene expression of selected genes as in Figure 1B of each cluster. **(H)** The thresholds of identifying CD45.1+ and CD45.2+ cells in day0, day1, day7, day14, day56 STIA CD45.1àCD45.2 BMC mice CITE-seq datasets. **(I)** Correlation of 8 module gene in merged day0, day1, day7, day14, day56 STIA CD45.1àCD45.2 BMC datasets. **(J)** Sankey plot of the contribution of 3 up- and down-regulated gene patterns identified in STIA B6 bulk RNA-seq dataset (Figure 3D) from 8 module scores identified in merged day0, day1, day7, day14, day56 STIA CD45.1‘*)CD45.2 BMC datasets. **(K)** Selected gene expression of 4 macrophage subpopulations stratified by timepoint.

**Supplemental Figure 6.**
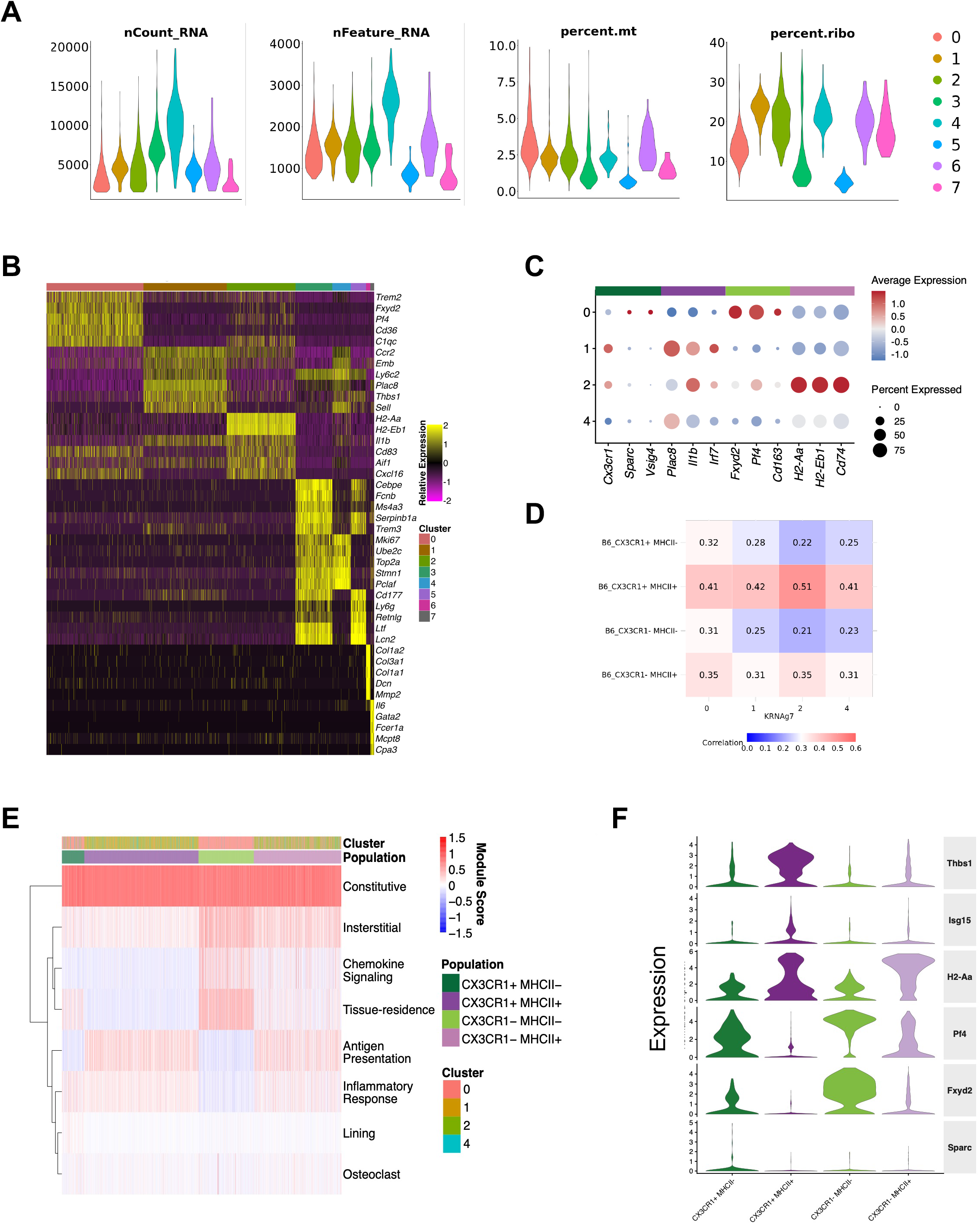
**(A)** CITE-seq QC results of nCount_RNA, nFeature_RNA, percent.mt, percent.ribo, nCount_ADT of each cluster in KRNAg7 dataset. **(B)** Heatmap of representative gene markers for each cluster. **(C)** Bubble plot gene expression of selected genes as in Figure 1B of each cluster. **(D)** Correlation between B6 macrophage subpopulations and 4 clusters in KRNAg7 CITE-seq dataset. **(E)** Module score heatmap of 9 module genes in each population stratified by cluster. **(F)** Expression of selected gene markers across 4 macrophage subpopulations.

## Notes

### Competing Interest Statement

The authors have declared no competing interest.

### Funding

- NIH: R01 AI163742, R01 AR080513
- Arthritis National Research Foundation: Hevolution/AFAR

